# Toxin-like neuropeptides in the sea anemone *Nematostella* unravel recruitment from the nervous system to venom

**DOI:** 10.1101/2020.05.28.121442

**Authors:** Maria Y. Sachkova, Morani Landau, Joachim M. Surm, Jason Macrander, Shir Singer, Adam M. Reitzel, Yehu Moran

## Abstract

The sea anemone *Nematostella vectensis* (Anthozoa, Cnidaria) is a powerful model system for characterizing the evolution of genes functioning in venom and nervous systems. Despite being an example for evolutionary novelty, the evolutionary origin of most toxins remains unknown. Here we report the first bona fide case of protein recruitment from the cnidarian nervous to venom system. The ShK-like1 peptide has ShKT cysteine motif, is lethal for fish larvae and packaged into nematocysts, the cnidarian venom-producing stinging capsules. Thus, ShK-like1 is a toxic venom component. Its paralog, ShK-like2, is a neuropeptide localized to neurons and is involved in development. Interestingly, both peptides exhibit similarities in their functional activities: both of them provoke contraction in *Nematostella* polyps and are toxic to fish. Because ShK-like2 but not ShK-like1 is conserved throughout sea anemone phylogeny, we conclude that the two paralogs originated due to a *Nematostella*-specific duplication of a ShK-like2 ancestor, a neuropeptide-encoding gene, followed by diversification and partial functional specialization. Strikingly, ShK-like2 is represented by two gene isoforms controlled by alternative promoters conferring regulatory flexibility throughout development. Additionally, we characterized the expression patterns of four other peptides with structural similarities to studied venom components, and revealed their unexpected neuronal localization. Thus, we employed genomics, transcriptomics and functional approaches to reveal one new venom component, five neuropeptides with two different cysteine motifs and an evolutionary pathway from nervous to venom system in Cnidaria.

## Introduction

The starlet sea anemone *Nematostella vectensis* has developed into a model system for characterizing the evolution of genes and their functions. It has a non-centralized nervous system, which comprises morphologically distinguishable sensory, ganglion and nematocyte neuronal types that develop from the same neural progenitor cells (1). From a transcriptomic perspective, however, several dozen neuronal subtypes were identified (2). Unlike in bilaterian animals where neurons differentiate from and are located in the ectoderm, the nervous system of *Nematostella* spans both ectoderm and endoderm (3, 4). Ganglion cells are found in both germ layers while sensory cells and nematocytes are present only in the ectoderm. Nematocytes are highly specialized cells bearing a nematocyst, an organelle loaded with proteinaceous venom that can be discharged upon mechanical stimulation (5-8). Thus, nematocytes function as micro-syringes injecting venom into prey and predators.

Sea anemone venom is a complex mixture of toxins. Along with nematocytes, ectodermal gland cells contribute to venom production (9, 10). A significant portion of the venom consists of disulphide-rich peptides of approximately 4 kDa molecular weight (11). Many of these peptides possess neurotoxic activity and act through binding to a target receptor in the nervous system of the prey or predator interfering with transmission of electric impulses. For example, Nv1 toxin from *Nematostella* inhibits inactivation of arthropod sodium channels (12) while ShK toxin from *Stichodactyla helianthus* is a potassium channel blocker (13). *Nematostella*’s nematocytes produce multiple toxins with a 6-cysteine pattern of the ShK toxin (9). Comparative phylogenetic analyses suggested that many venom components originated through recruitment of proteins from other physiological systems such as the digestive, immune and nervous systems (14). However, the origin for many animal toxins, particularly understudied groups like cnidarians, remains relatively unknown.

Cnidarian neurons produce multiple neuropeptides (4, 15, 16) that act as neurotransmitters and neuromodulators. Thirty putative neuropeptide sequences belonging to families widely represented in metazoans (such as RFamide, vasopressin, galanin, tachykinin peptides) as well as cnidarian-specific families (for example, Antho-RIamide, RNamide, RPamide peptides) have been identified in *Nematostella* genome (17). All the reported mature sequences are relatively short (3-15 residues) with a C-terminal amidation signal (Gly residue) and absence of Cys residues (apart of two RPamide peptides). Recently, an approach based on tandem mass spectrometry and a novel bioinformatics pipeline identified 20 new neuropeptide sequences in *Nematostella* (18). Some of the new peptides were classified as RFamide and RPamide peptides, which were reported earlier, while others did not share similarity to any known cnidarian neuropeptides.

Cnidarian neuropeptides affect diverse physiological processes such as muscle contraction, larval migration, metamorphosis, and neural differentiation (15, 16). *Nematostella* GLWamide neuropeptide regulates timing of transition from planula to polyp (19). Most of the studied metazoan neuropeptides act through binding to metabotropic receptors, such as peptide gated G-protein-coupled receptors (GPCRs) (20), however, RFamides from another cnidarian (*Hydra*) directly activate ion channels from the degenerin /epithelial Na^+^ channel family (4).

Venom components and neuropeptides are secreted molecules. Typically, secreted peptides are synthesized as precursor proteins equipped with a signal peptide leading to co-translational translocation of the precursor into the endoplasmic reticulum (21). In cnidarians, the precursor also typically includes a propeptide that is cleaved off by maturation enzymes upon processing (22). Mature peptides are transported to their secretion sites, which can be a neuronal synapse in case of neuropeptides or a nematocyst in case of venom components.

Here we report two novel classes of neuropeptides with structural similarity to venom components. We have characterized the novel paralogs expressed in venom and nervous systems at the genomic, transcriptomic and functional levels. Our work reveals a first case of protein recruitment from the nervous to venom system.

## Results

### New toxin-like peptides

The *Nematostella* transcriptome remains incompletely annotated, therefore, we expanded our search to identify all potential toxins from this species. We used toxin sequences deposited in UniProt (ToxProt) as a query to identify toxin-like sequences in *Nematostella* using the previously published TSA dataset from mesenteries, nematosomes, and tentacles (PRJEB13676). This search identified Class8-like peptide, a homolog of Avd8e toxin (e-value 7×10^−18^), and Aeq5-like1, a homolog of Acrorhagin 5a from *Actinia equina* (e-value 0.005) (23). Using Transdecoder (62), we translated the TSA sequences and obtained 1,126,514 protein sequences, with 653 sequences with the predictive ShK-like cysteine arrangement, 74 of which were predicted to have a signal region. After screening for multiple ShK-like domains, excessive cysteine residues, and sorting through ToxProt BLAST outputs, we identified transcripts encoding six uncharacterized proteins with a cysteine arrangement typical of the potassium channel blocker from *Stichodactyla helianthus* known as ShK (ShK-like1, ShK-like2a, ShK-like2b, ShK-like3, ShK-like4, and Class8-like; **Table 1, Fig 1A**). Additionally, we examined the *Nematostella* single cell RNAseq data (2)) where one gene, Nve8041, was expressed in neurons and coded for a protein with the same Cys pattern as Aeq5a and Aeq5-like1; that we designated as Aeq5-like2 (**Table 1, Fig 1B**).

**Table 1.** New toxin-like peptides. Gene models are according to Fredman D et al 2013, https://doi.org/10.6084/m9.figshare.807696.v1

| Name | TSA ID/ Nve gene model | Homolog in ToxProt | Activity of the ToxProt homolog | % identity (ClustalO) | Cys motif |
| --- | --- | --- | --- | --- | --- |
| ShK-like1 | HADP01047794.1/NA | *A. viridis* Avd9a | Toxin by similarity | 30% | ShKT |
| ShK-like2a | HADN01006514.1/NA | *A. viridis* Avd9a | Toxin by similarity | 24% |  |
| ShK-like2b | HADO01001456.1/Nve11386 | *A. viridis* Avd9a | Toxin by similarity | 25% |  |
| ShK-like3 | HADN01040061.1/NA | *A. viridis* Avd9c | Toxin by similarity | 34% |  |
| ShK-like4 | HADO01004651.1/Nve12327 | *A. viridis* Avd9b | Toxin by similarity | 32% |  |
| Class8-like | HADO01000486.1/Nve7851 | *A. viridis* Avd8e | Toxin by similarity | 51% |  |
| Aeq5-like1 | HADO01002561.1/Nve8064 | *A. equina* Aeq5a | Extracted from acrorhagi, toxic for crabs | 39% | Aeq5a |
| Aeq5-like2 | Nve8041 | - | - | - |  |

**Figure 1.**
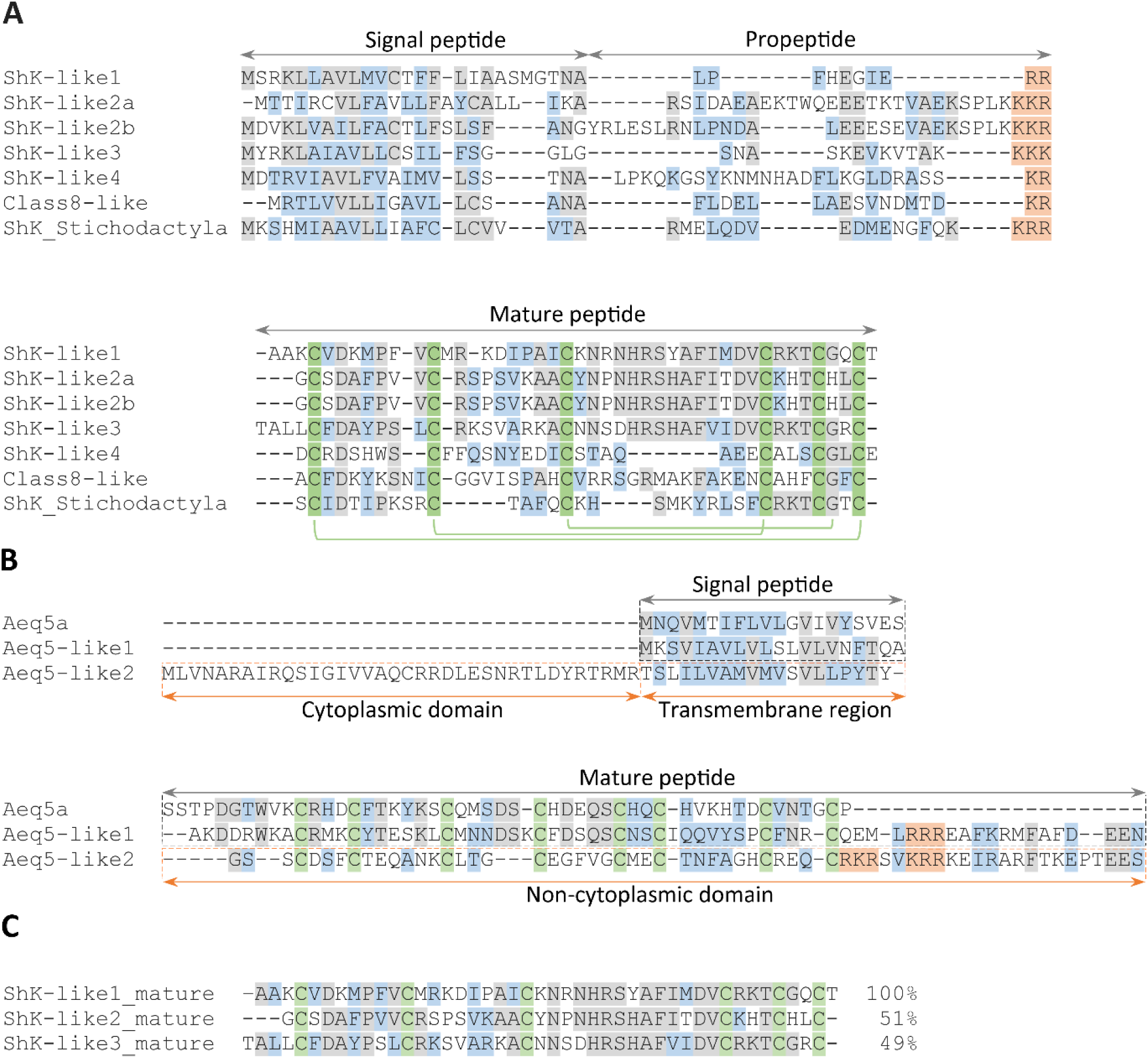
Alignment of the new toxin-like precursor sequences from Nematostella and characterized toxins ShKT from Stichodactyla helianthus (A) and Aeq5a from Actinia equina (B). The new sequences were split into signal peptide, propeptide and mature peptide fragments and corresponding fragments were aligned separately using Muscle algorithm implemented in MegaX and manually adjusted. Processing sites are in orange, cysteine residues making part of the cysteine motifs of the mature peptides are in green. Identical residues are in grey, conserved substitutions are in blue. (**A**) Sequences with ShKT cysteine motif, disulphide bonds of ShK toxin are shown as green brackets. **(B)** Sequences with the same motif as in Aeq5a. **(C)** Alignment of the ShK-like1, ShK-like2 and ShK-like3 mature peptides.

All the newly identified proteins, with the exception of Aeq5-like2, possess structural features typical of sea anemone toxin precursor proteins: a signal peptide (identified by SignalP (24)), a propeptide with a dibasic processing motif (apart of Aeq5-like1) followed by a Cys rich mature peptide (25). ShK-like2a and ShK-like2b differ in signal peptides and a portion of the propeptides, while their mature peptides are identical (therefore referred to as ShK-like2). ShK-like1, ShK-like2a/b, and Shk-like3 are likely closely related homologs as their mature peptides share around 50% identical residues (**Fig 1C**). Signal peptides and propeptides are more diverged as their overall identity level is 36% to 40%. Aeq5-like2 has slightly different structure – instead of a signal peptide it has a transmembrane domain on its N-terminus indicating that the disulphide-rich domain is anchored on the extracellular side of the cell membrane (InterProScan prediction (26)). Furthermore, the validity of these newly identified transcripts as protein-coding genes was confirmed at the protein level with tandem mass spectrometry (LC-MS/MS) data (9) detecting ShK-like1, ShK-like2, ShK-like4 and Aeq5-like1 at multiple life-stages in *Nematostella* (**Suppl fig 1**).

ShK-like1 biosynthesis dynamics differed from the other proteins: it was the most abundant in larvae and significantly decreased later in the life cycle, according to transcriptomic (27) and proteomic (9) data (**Suppl fig 1**). On the contrary, all the other genes apart of ShK-like3 were expressed early in the life cycle and maintained expression through the adult stage. For ShK-like3, however, no reads mapped to the transcript in the NveRtx database (ID: NvERTx.4.119933).

### ShK-like1 is a venom component

We used whole mount in situ hybridization (ISH) in *Nematostella* planulae and primary polyps to determine the expression patterns of the newly discovered toxin-like genes. Surprisingly, only ShK-like1 was expressed in nematocytes of planulae and primary polyps (**Fig 2A**). At the transcriptomic level (NvERtx database (27)), ShK-like1 expression in planulae was comparable to the previously characterized Nep3 toxin (9). In the primary polyps, ShK-like1 was localized predominantly in the body column nematocytes and not in the tentacles.

**Figure 2.**
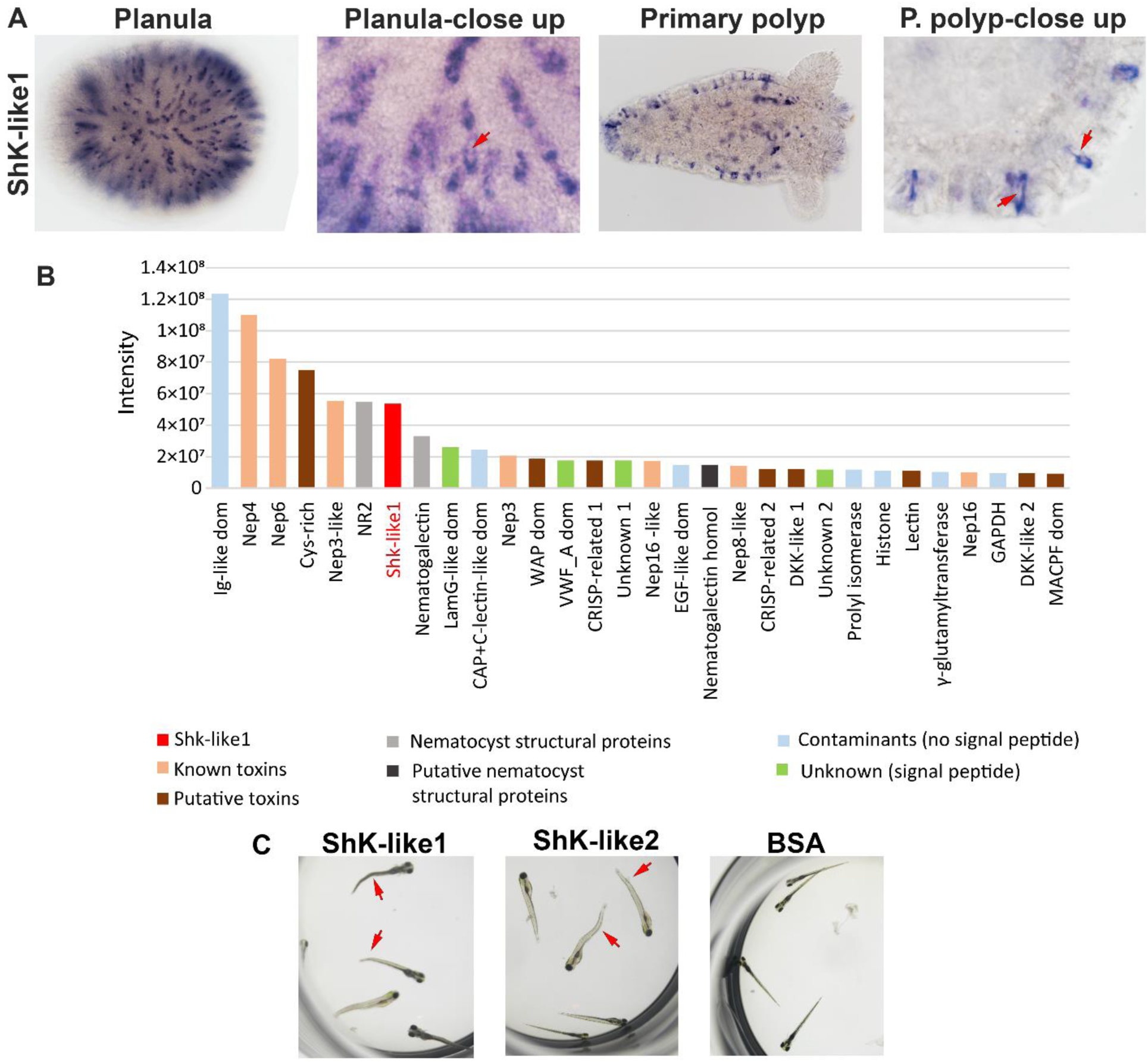
ShK-like1 is a venom component. (**A**) ShK-like1 gene is expressed in nematocytes in planula and primary polyps; note that the staining is distributed to the periphery of the cells and the unstained center of the cells corresponds to capsules (arrows). (**B**) 30 most abundant proteins extracted from planula nematocysts identified by tandem mass spectrometry. (**C**) Toxicity of recombinant ShK-like1 and ShK-like2 was tested on zebrafish larvae in parallel to BSA control. Arrows point to twitched tails evidencing for the contraction paralysis effect of the peptides.

To confirm that ShK-like1 protein is packaged into nematocysts and thus is a venom component, we analyzed soluble protein content of nematocysts by LC-MS/MS. Because abundance of the ShK-like1 protein is higher in larvae than in adults by more than one order of magnitude (**Suppl fig 1D**), we used capsules from 4 days post-fertilization (dpf) planulae. 138 proteins with at least two unique tryptic peptides were identified. ShK-like1 (4 unique peptides) along with other previously characterized nematocyst toxins (Nep3, Nep3-like, Nep4, Nep6, Nep8-like, Nep16 (7, 9)) and nematocyst structural proteins were among the 30 most abundant proteins (**Fig 2B, Suppl table 1**).

To study the toxic effects of ShK-like1, the mature peptide was produced as a recombinant protein and tested on zebrafish larvae. After 1 h incubation with 0.5 mg/ml ShK-like1, 20 out of 20 larvae started exhibiting contraction paralysis and within 2 h they died while all the larvae in BSA control survived (**Fig 2C**). That evidences for neurotoxic activity of ShK-like1. Thus, ShK-like1 is a nematocyst toxin highly expressed at the planula.

### The unexpected expression of toxin-like genes in neurons

In *Nematostella*, ganglion and sensory neurons can be recognized by their morphological features: ganglion neurons have 2-4 neural processes while sensory cells have elongated shape, possess processes at the base and may have a cilium at the apex (28). Unexpectedly, ISH showed that ShK-like2a/b, ShK-like4, Class8-like, Aeq5-like1, and Aeq5-like2 were localized in various neurons in both planula and primary polyp stages (**Fig 3-7**).

**Figure 3.**
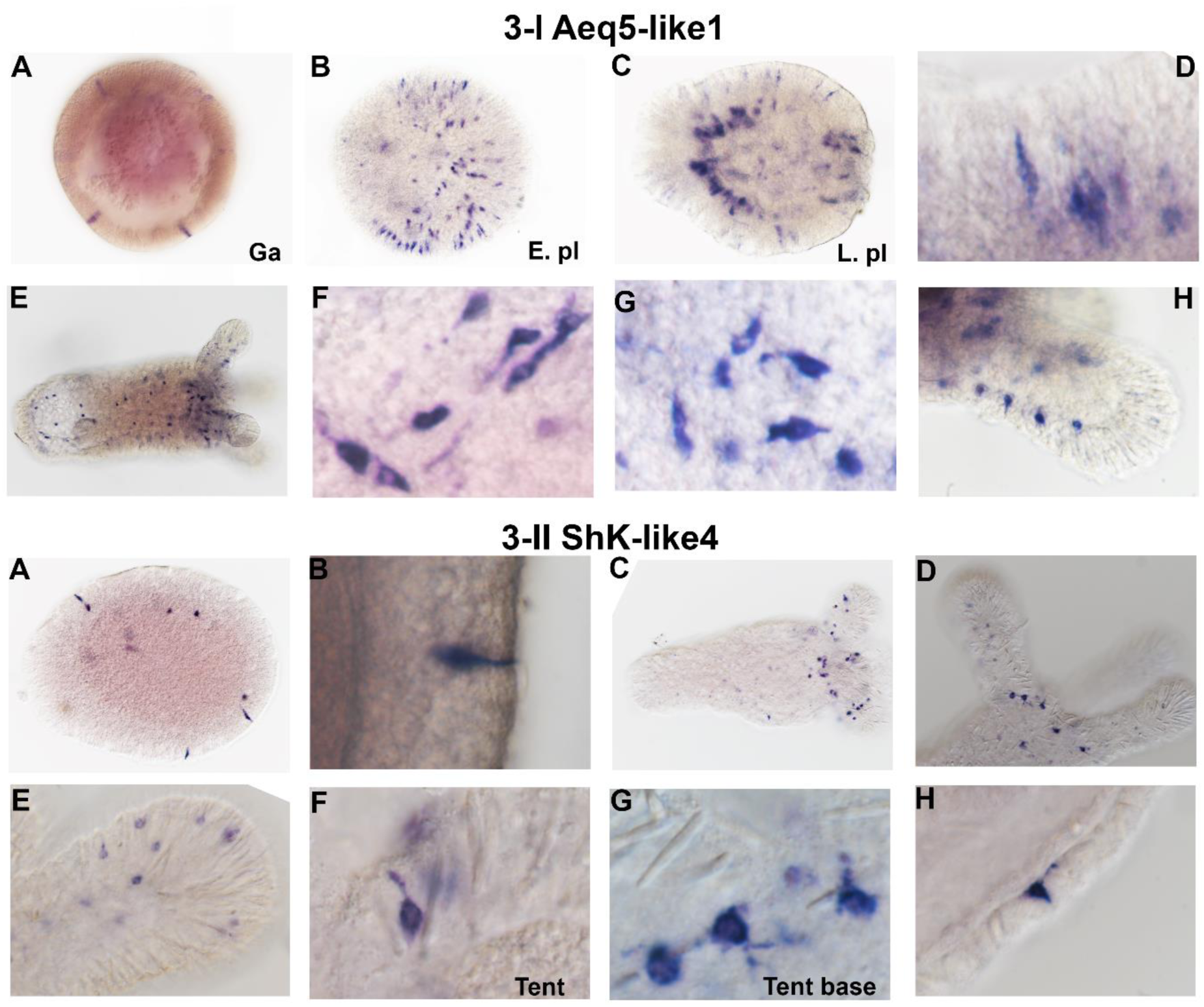
Expression pattern of Aeq5-like1 and ShK-like4 genes by ISH. 3-I: Aeq5-like1 expression starts in the ectodermal cells of gastrula (**A**) and early planula (**B**) and by the late planula (**C**) it is also noticeable in the endoderm. (**D**) Morphology of the planula ectodermal cells is consistent with sensory neurons. In the primary polyp, Aeq5-like1 is expressed in both ectoderm and endoderm (**E**) in ganglion (**F, G**) and sensory neurons (**H**). 3-II: ShK-like4 gene is expressed in ectodermal sensory cells in planula (**A, B**) and primary polyp (**C** - **H**).

Aeq5-like1 expression was first detected early in development in ectodermal cells of the gastrula, then increased in early planula and by the late planula had spread in expression to endodermal cells **(Fig 3-I)**. Early ectodermal cells may represent neuronal precursors for sensory cells. In primary polyps, Aeq5-like1 was expressed in ectodermal sensory cells and multiple endodermal ganglion cells that send their processes in different directions all around the body. ShK-like4 was expressed in ectodermal sensory cells in planulae and oral part of polyps **(Fig 3-II)**. Some of the sensory cells appeared to have cilia (**Fig 3-II B, F**) and processes at their base (**Fig 3- II G, H**). Class8-like was expressed in multipolar ganglion cells with long branching processes localized in mesoglea (**Fig 4-I**). Aeq5-like2 was localized to endodermal ganglion neurons, apparently bipolar and following mesentery folds (**Fig 4-II**). To the best of our knowledge these patterns are quite unique as in most cases, unlike protein-based staining, RNA-based staining of *Nematostella* neurons does not reveal neurite like structures (28, 29).

**Figure 4.**
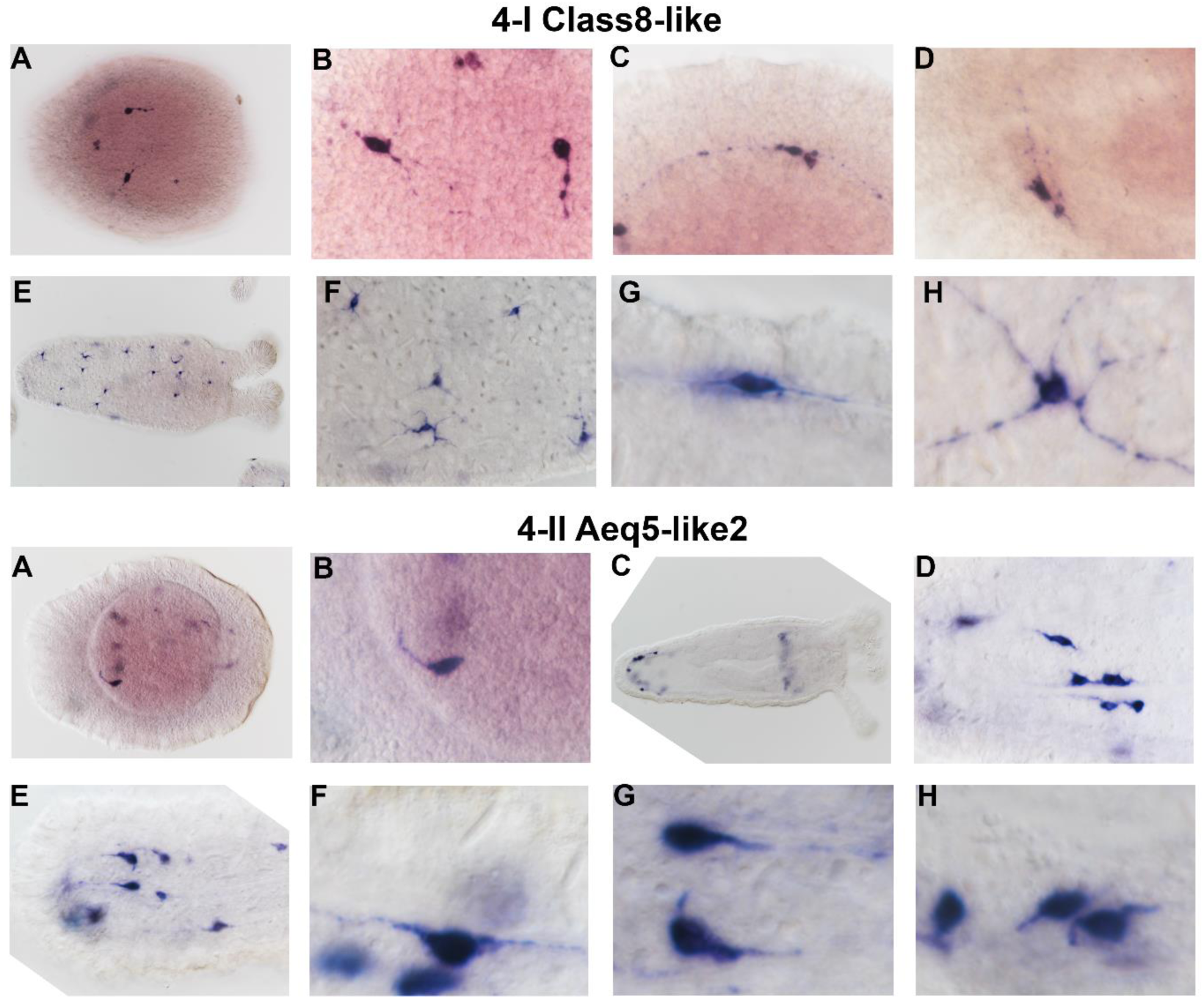
Expression pattern of Class8-like and Aeq5-like2 genes by ISH. 4-I: Class8-like gene is expressed in the ganglion neurons residing in the mesoglea of planula (**A-D**) and primary polyp (**E-H**). **4-II:** Aeq5-like2 is expressed in the endodermal ganglion neurons in planula (**A**, **B**) and primary polyp (**C - H**).

For ShK-like2 ISH, at first we synthesized an RNA probe that was relatively long (ShK-like2-long, 603 nt) and specific to the full-length ShK-like2b, which also shared significant portion of the sequence (66%) with the ShK-like2a mRNA (**Suppl table 2, Fig 5-I**). The resulting pattern allowed us to conclude that ShK-like2 was predominantly expressed in ectodermal sensory neurons in planulae and endodermal ganglion cells in primary polyps. To determine whether ShK-like2a and ShK-like2b exhibit different expression patterns, we produced probes that were specific for each isoform. Due to relatively short coding sequences, ShK-like2a probe (ShK-like2a-short, 372 nt) included 84% of specific sequence while ShK-like2b probe (ShK-like2b-short, 332 nt) possessed 62% of specific sequence with the rest of the sequence shared between the two isoforms (**Suppl table 2**). ISH with the specific probes showed that ShK-like2b was first expressed in the early planula in ectodermal cells and this pattern persisted through the late planula stage, while ShK-like2a was first expressed mostly in the endoderm (with rare ectodermal cells) in the late planula. In primary polyps, expression patterns were indistinguishable: both isoforms were highly expressed in endodermal ganglion cells in the body column and tentacles with rare sensory ectodermal cells (**Fig 5-II, 5-III**). This pattern was confirmed by significant overlap between stained cells in double fluorescent in situ hybridization (dFISH), especially in the tentacle and aboral endoderm (**Suppl fig 2**). ISH with a probe for ShK-like3 transcript did not show any specific staining (**Suppl fig 3**).

**Figure 5.**
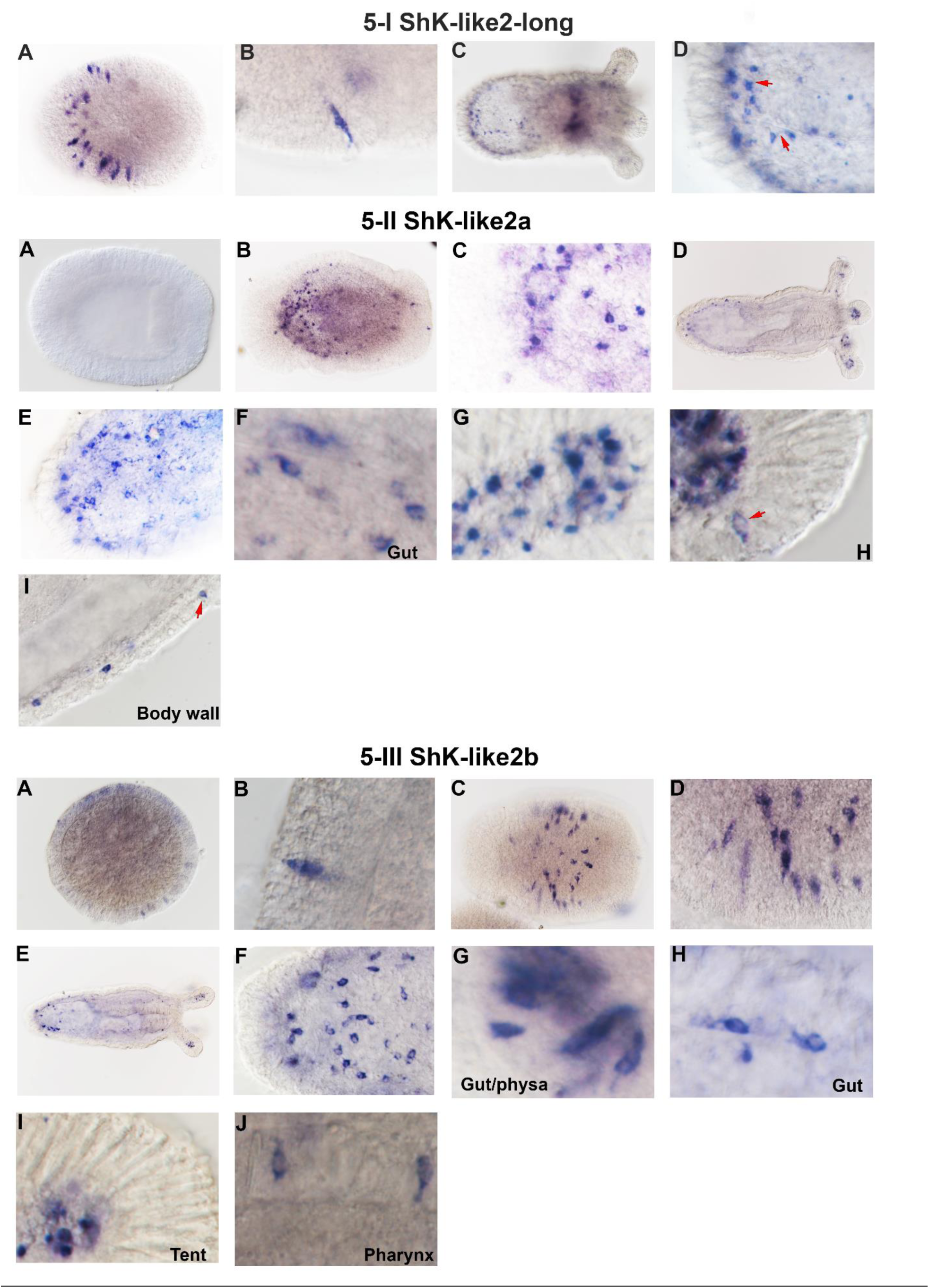
ShK-like2 expression pattern by ISH. (5-I) ISH with the ShK-like2-long probe reveals expression in planula’s ectodermal cells (**A**) with a sensory neuron morphology (**B**) and in primary polyp’s endoderm (**C**) in ganglion neurons with characteristic protrusions (**D**, arrows). **(5-II**) ISH with ShK-like2a-short probe reveals that ShK-like2a has no expression in early planula (**A**), expression starts mostly in the endoderm of late planula (**B**) in ganglion neurons (**C**). In the primary polyp (**D**), it is expressed mostly in the endodermal ganglion neurons in physa (**E**), gut (**F**) and tentacles (**G, H**). The expression is also evident in the smaller number of ectodermal sensory neurons in tentacles (**H**, arrow) and body wall (**I**, arrow). (**5-III**) ISH with ShK-like2b-short probe shows that ShK-like2b starts expressing in the ectoderm of early planula (**A**) in elongated cells (**B**) and continues in the ectodermal sensory neurons in the late planula (**C, D**). In the primary polyp (**E**), it is expressed in multiple endodermal ganglion neurons (**F**) in physa (**G**), gut (**H**) and tentacles (**I**). ShK-like-2b is also found in a small number of ectodermal neurons in pharynx (**J**).

To determine whether the novel neuropeptides are co-localized with the well-studied neuronal marker ELAV (3), we employed double in situ hybridization (dISH) in primary polyps. ShK-like2 expression (the long probe) showed partial overlap with ELAV in tentacle endoderm but not in the aboral endoderm. ShK-like4 expression partially co-localized with ELAV in the ectodermal sensory cells. Aeq5-like1 and Class8-like expression did not show any overlap (**Suppl fig 4-5**).

Single cell RNAseq data (2) confirmed that ShK-like2b, ShK-like 4, Aeq5-like1, Class8-like and Aeq5-like2 are expressed in neurons in both larvae and adult polyps (**Suppl fig 6**). In larvae, ShK-like2b and Class8-like expression were additionally upregulated in undifferentiated metacells that may correspond to neural precursors. In adult polyps, these data also supported partial overlap between ELAV and the neuropeptide expression that we observed in dISH on primary polyps: ShK-like4 in the C32 neuronal metacell and ShK-like2b in C34. It also suggested an overlap of Aeq5-like1 and ELAV in C30 neurons that we were not able to detect in our dISH experiments.

**Figure 6.**
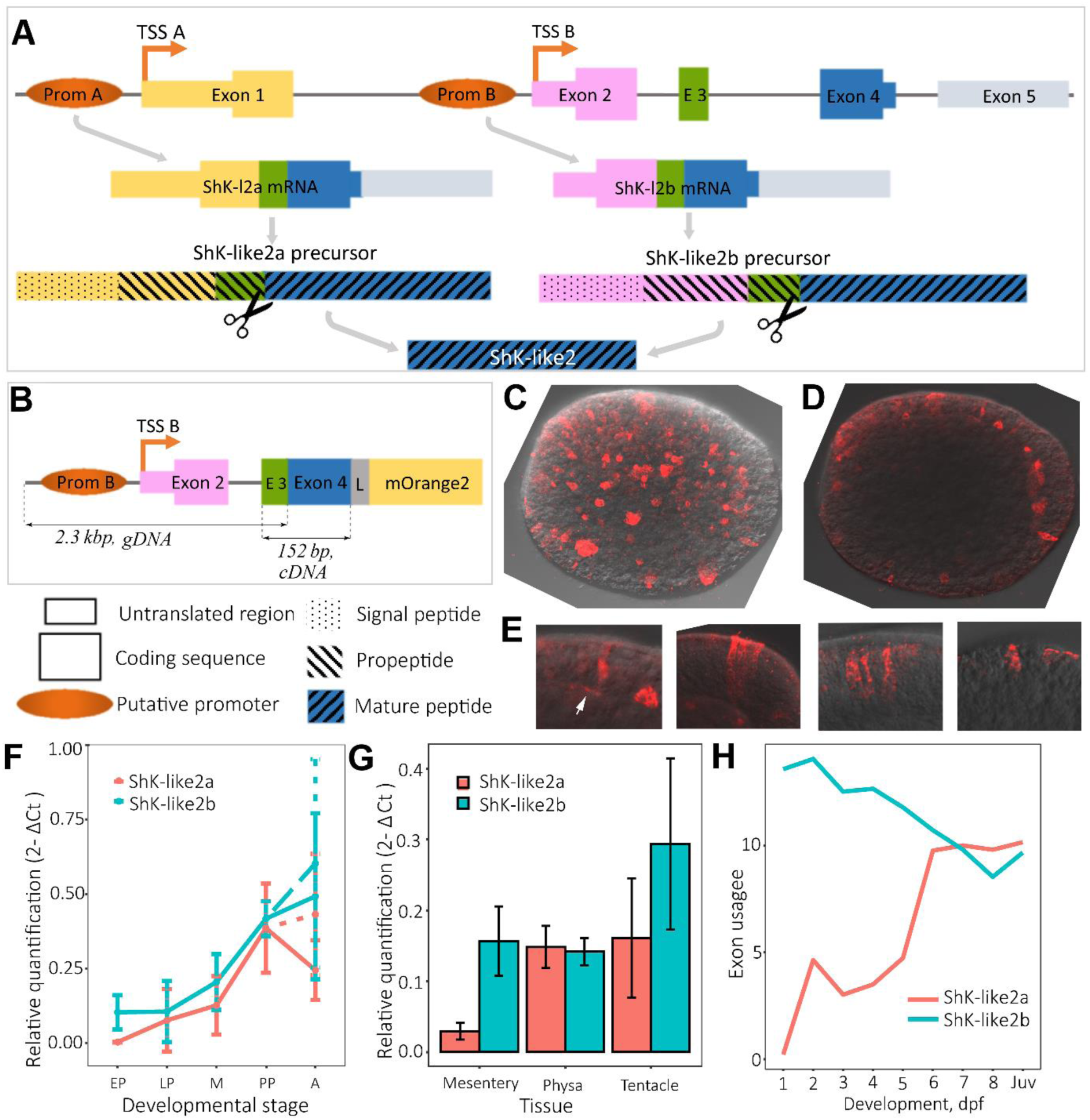
Structure and regulation of ShK-like2 gene. **(A)** ShK-like2 gene structure and biosynthesis of the two isoforms of ShK-like2 precursor protein leading to the same mature peptide. (**B**) Scheme of the transgenic construct aimed to label cells expressing ShK-like2b isoform. (**C - E**) Immunostaining of transgenic planulae, confocal imaging; red – mOrange2. Staining in multiple ectodermal cells (Z-stack of 42 maximum intensity projections). (**D**) Individual stained ectodermal cells (Z-stack of 4 maximum intensity projections). (**E**) Stained cells are morphologically sensory neurons with processes at the base (arrow) and uneven distribution of the staining evidencing for mOrange2 being packed in vesicles (Z-stacks of 3 to 4 maximum intensity projections). (**F**) ShK-like2a and ShK-like2b gene expression dynamics in development measured by qPCR. EP – early planula, LP – late planula, M **–** metamorphosis, PP – primary polyp, A – adult; adult females – solid line, adult males – dashed line. (**G**) ShK-like2a and ShK-like2b expression levels in adult tissues measured by qPCR. In **F** and **G**, error bars represent standard deviations. (**H**) Differential usage of exon 1 (ShK-like2a) compared to exon 2 (ShK-like2b) detected by DEXseq. Juv – juvenile polyp (6 weeks old).

### ShK-like2 gene structure and evolution

Alignment of the ShK-like2a and ShK-like2b transcripts to the genome (30) showed that both isoforms were encoded by the same gene (scaffold_260:199,854-210,905) with alternative transcription start sites (TSS) (**Fig 6A**). The ShK-like2 gene contains four coding exons, where exon 1 and exon 2 are alternating between the isoforms. Transcription starting from putative promoter A leads to ShK-like2a encoded by the exons 1, 3, and 4 while alternative transcription activated by putative promoter B results in ShK-like2b encoded by the exons 2, 3, and 4. To confirm that ShK-like2b expression was driven by the alternative promoter B localized upstream of exon 2, we generated a transgenic construct bearing the reporter gene mOrange2 (31) under the control of ShK-like2b regulatory elements. Injection of the transgenic construct into *Nematostella* zygotes resulted in reporter gene expression in ectodermal cells of the planulae (**Fig 6B-E**).

RT-qPCR analysis showed that ShK-like2 was initially expressed either in early planula (ShK-like2b isoform) or late planula (ShK-like2a isoform) and increased to the adult stage, with the exception of ShK-like2a in adult females that appeared to decrease its expression level (**Fig 6F)**. Both isoforms were expressed in the physa, mesentery and tentacles of adult females; ShK-like2b had higher expression level in the mesentery (**Fig 6G)**.

To confirm the differences between the expression profiles of ShK-like2a and ShK-like2b isoforms observed in the qPCR and ISH staining (**Fig 5-II, 5-III**), we employed DEXseq (32). Raw reads, generated from *Nematostella* across a developmental time course (1-8 dpf and juveniles) (27), were mapped back to a *de novo* assembled transcriptome to detect differential exon usage. DEXseq analysis confirmed that exon 1 (ShK-like2a) and exon 2 (ShK-like2b) had significant differential exon usage (FDR p-value < 0.001) across development. Specifically, ShK-like2b was preferentially expressed from gastrula stage (1 dpf) to late planulae stage (5 dpf), with exon 2 usage having a fold change ≥ 2 compared to exon 1. Following metamorphosis (6 dpf), no significant difference in exon usage was observed (**Fig 6H**). Thus, DEXseq analysis detected that patterns of ShK-like2 exon expression were consistent with our qPCR results.

Interestingly, structure of the genomic locus of the ShK-like2 orthologue is conserved in the sea anemones *Exaiptasia pallida* and *A. equina*, which are distantly related to *Nematostella* (33, 34). In these species, regulation of the ShK-like2 gene by alternative TSSs also results in a protein with altering signal peptides (35). ShK-like2 orthologues retrieved by BLAST search in other available sea anemone transcriptomes (*Calliactis polypus, Nemanthus annamensis, Aulactinia veratra, Actinia tenebrosa, A. equina (36), Anthopleura buddemeieri* (37), *Anthopleura elegantissima, Edwardsiella lineata* (38)) also possessed alternative signal peptides and identical mature parts (**Fig 7A, B)**, suggesting the regulatory mechanism is widely conserved in sea anemones. Thus, it is likely that a ShK-like2 orthologue with this unique structure and regulation was present in the last common ancestor of all sea anemones.

**Figure 7.**
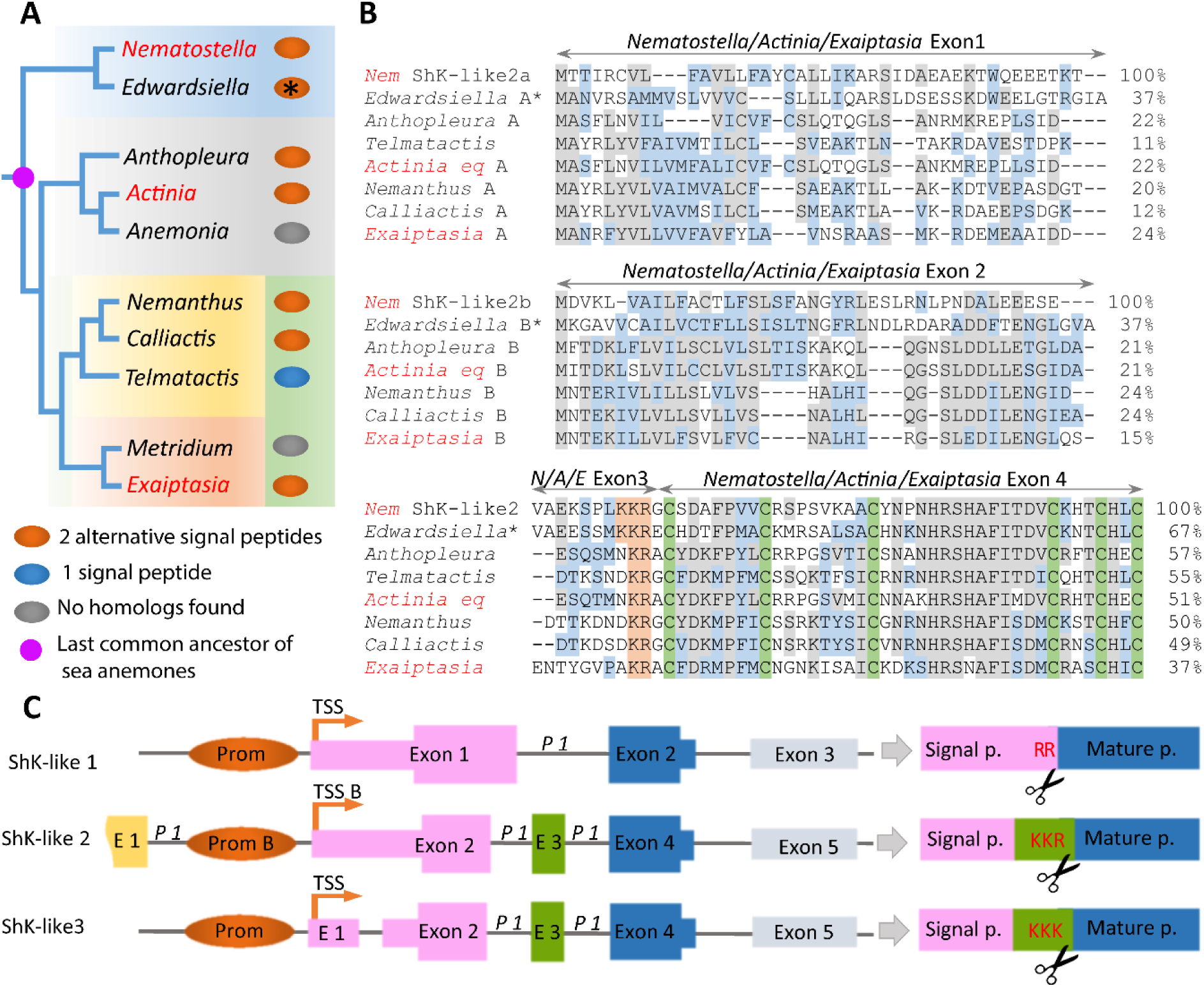
Evolution of ShK-like2 gene. **(A)** Distribution of the ShK-like2 homologs across the phylogenetic tree of sea anemones; the species with sequences genomes are in red (33). The number of alternative signal peptides for each ShK-like2 homolog identified in each genus (transcriptome or genome) is represented by ovals (orange – 2, blue – 1, grey – no homologs); the asterisk denotes that full length sequences were obtained by joining shorter fragments manually. (**B**) Alignment of ShK-like2 homologs from different sea anemones. Exon-intron structure is conserved between Nematostella, Actinia and Exaiptasia (in red) and shown above the alignment. (**C**) Alignment of exon-intron structures of ShK-like1, ShK-like2 and ShK-like3 genes.

Gene models for ShK-like1 and ShK-like3 had not been established before, therefore we aligned corresponding transcripts to the genome scaffolds. ShK-like1 transcript aligned to a region of scaffold 53 which was interrupted by a gap corresponding to a part of the mature peptide. We PCR amplified and sequenced that fragment of the genomic DNA to annotate the remaining part of this locus (**Suppl table 3**). Structure of the ShK-like1 (scaffold_53:491,201-496,216) and ShK-like3 (scaffold _7:1,285,234-1,288,665) genes was quite different: ShK-like1 gene had 2 coding exons while ShK-like3 gene contained 3 coding exons (**Fig 7C**). The ShK-like3 gene structure was similar to ShK-like2 gene fragment encoding ShK-like2b. Orthologues of Shk-like1 and ShK-like3 were not found in other sea anemones, not even in other members of the Edwardsiidae family. Thus, ShK-like1 and ShK-like3 probably resulted from *Nematostella*-specific gene duplications followed by sequence divergence. ShK-like3 is expressed at very low level and ISH did not result in any staining therefore it is probably a pseudogene.

### Functional characterization of ShK-like2

Because ShK-like2 shares high sequence similarity with the ShK-like1 toxin we were interested in characterizing its biological activity in comparison to ShK-like1. To determine whether ShK-like2 provokes any noticeable physiological reaction in primary polyps, we incubated 11 dpf polyps with recombinant ShK-like2 in parallel to ShK-like1 and NvePtx1 toxins (0.5 mg/ml) in addition to the BSA control (5 mg/ml). After 2 h, length to width ratio of the body column was significantly lower in the ShK-like2 sample compared to NvePtx1 and BSA (p=2.4×10^3^ and p=3.4×10^8^, respectively; Student’s T-test) due to body contraction; there was no difference in comparison to the ShK-like1 sample (p=0.3, Student’s T-test) (**Fig 8A, B**).

**Figure 8.**
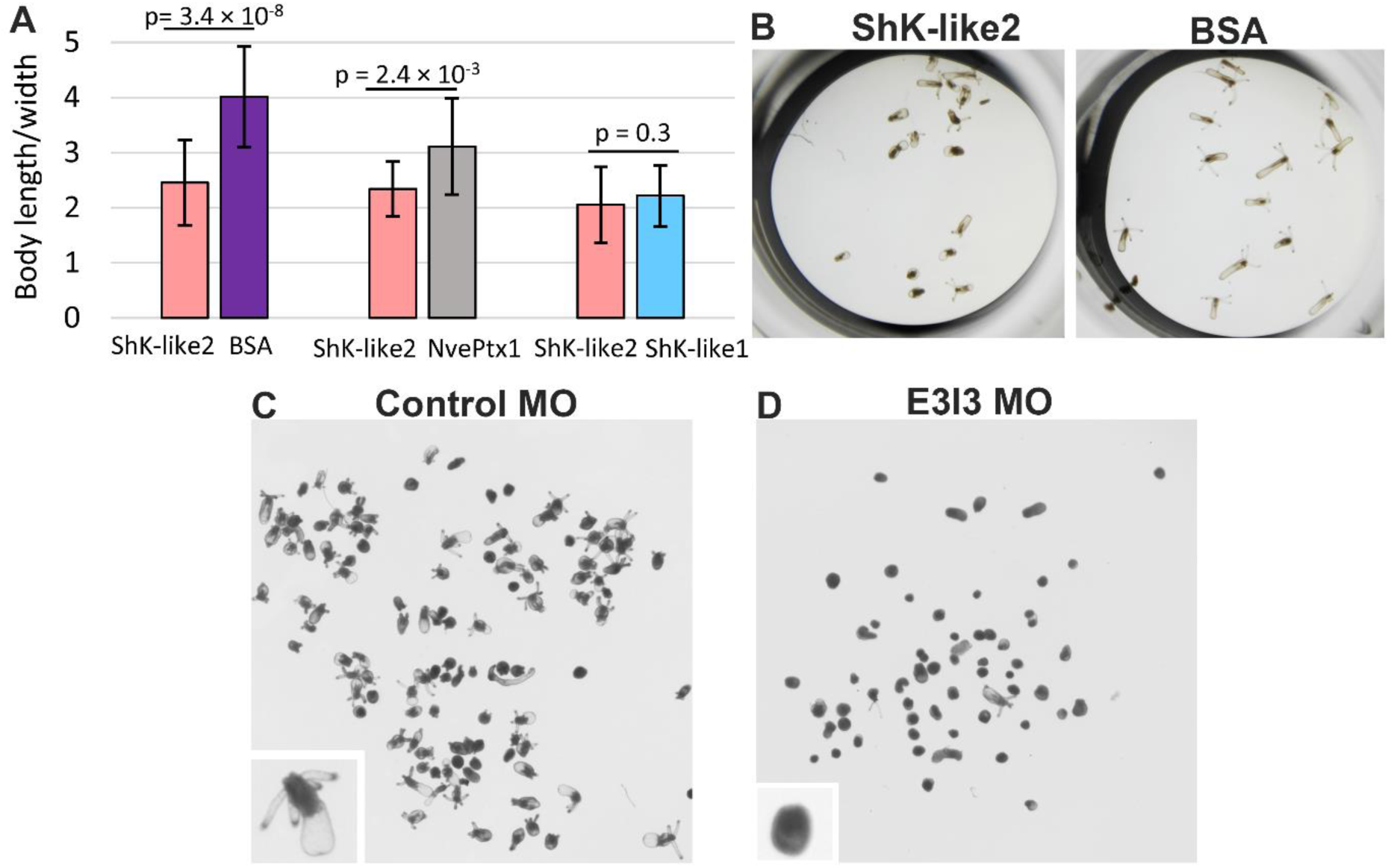
Effects of ShK-like2 on Nematostella. **(A)** Ratios of body length to width in primary polyps after 2 h of incubation with ShK-like2 in comparison to BSA, NvePtx1 and ShK-like1. (**B**) Effect of the 2 h incubation with ShK-like2 in comparison to BSA. (**C**-**E**) Developmental effects of MOs in 11 dpf polyps; close ups to embryos with typical phenotypes are in the lower left corners. (**C**) Control MO, (**D**) E3I3 MO.

To test the toxic activity of ShK-like2, we incubated zebrafish larvae in 0.5 mg/ml ShK-like2 solution in parallel to ShK-like1 and BSA control (**Fig 2C**). We observed paralysis and death in 4 out of 15 tested larvae within 2 h of incubation.

ShK-like2 was first expressed at the early planula stage and therefore we aimed to test whether it played a role in development using gene knockdown with Morpholino oligonucleotides (MO). We aimed to disrupt splicing for either exon 3 or exon 4, which are shared among the isoforms (**Fig 8A**), using specific MO sequences. Animals injected by either MO targeting exon 3 (E3I3) or 4 (I3E4) had strong developmental defects (**Fig 8C-D**), resulting in a significant proportion of larvae being unable to settle compared to a morpholino control (p=0, Chi-Square test and p < 0.00001, Fisher exact test for E3I3 and I3E4, respectively). In comparison to the control MO, injection of E3I3 and I3E4 MOs on average reduced planulae settlement by 39.4±8.8% (4 replicates, **Suppl table 5**) and 83.4±9.4% (3 replicates, **Suppl table 6**), respectively. The difference in the effects may be explained by the difference in efficiency of each MO. PCR analysis showed that I3E4 led to nearly complete elimination of the processed mRNA, while E3I3 MO injection resulted in partial reduction of the processed mRNA and correctly processed mRNA could be detected as well (**Suppl fig 7**).

## Discussion

### Annotation of toxins in cnidarians

The research reported here led to the discovery of the two novel neuropeptide types; however, it was initially aiming to find new toxins in *Nematostella* transcriptome using UniProt sequences annotated as toxins (including “toxin by similarity”) and search for secreted peptides with ShKT motif. Unnexpectedly, all but one of the retrieved candidates were found to be expressed in ganglion and sensory neurons, not in nematocytes or ectodermal gland cells. This demonstrates that molecules with structural similarities to venom components have wide expression in different cell types in cnidarians, particularly neurons. Thus, many sequences annotated as venom components based on the sequence similarity or toxic activity without dissecting their tissue localization may have different functions. This mischaracterization is a challenge to overcome as the cnidarian venom system composed of sparse, dispersed cells rather than anatomically defined venom gland leading to technical difficulties in studying venom-specific transcriptomes. Techniques enabling studies of cell-specific transcriptomes have only recently become available in cnidarians and will make this task more manageable (2, 39).

### Recruitment to venom

To the best of our knowledge, the recruitment of ShK-like1 is the first well-established case of a neuropeptide being recruited as a toxin. Our data provide evidence that ShK-like1, a newly discovered component of nematocyte venom, evolved by a *Nematostella*-specific duplication of a neuronally-expressed gene followed by sequence divergence and recruitment of one of the copies into the venom system. The other copy most probably retained its expression pattern giving rise to ShK-like2. ShK-like3 appears to result from a duplication of a ShK-like2 ancestral gene as well; however, unlike ShK-like1, it became a pseudogene. We have previously described a similar evolutionary pattern of gene gain and loss for another family of *Nematostella* venom components (40).

As neuropeptides affect the nervous system it might seem plausible they will be frequently recruited to venoms. However, such events were rarely described before, and it is possible that most neuropeptides make poor toxins because they act mainly via GPCRs, which deliver signal too slowly to be an effective target for toxins (20). An interesting example of short linear neuropeptides recruited to venom on multiple instances is invertebrate tachykinins (41). An evolutionary relationship between peptides from nervous and venom systems was also suggested for three-finger toxins (TFTx) in snakes (42, 43). However, the homology level between the TFTx of snakes and vertebrate neuropeptides is quite low and cannot be detected by BLAST.

### Partial functional specialization of ShK-like1 compared to ShK-like2

When we compared the toxicity of ShK-like1 and ShK-like2 on fish, it was noticeable that the latter is less effective, suggesting that ShK-like1 evolved higher pharmacological efficiency against fish. When new peptides are recruited into the venom system as toxins they may lose their original function. However, when we tested the ability of ShK-like1 to induce effects in *Nematostella* it was as effective as ShK-like2 (**Fig 10A**). Thus, it seems that ShK-like1 retained its ancestral activity to affect the *Nematostella* nervous system, despite the increased activity on fish. This suggests that increasing potency as a toxin against one species does not necessarily result in decreasing its efficacy on another. It is currently unknown what are the pharmacological targets or receptors of ShK-like1 and ShK-like2 and whether these targets are shared between fish and sea anemones.

### Cysteine-rich peptides in the nervous system and other regulatory systems

Secreted peptides with ShK-like cysteine motifs have been reported as venom components in sea anemones, however, to the best of our knowledge, the present work is the first evidence of ShK-like peptides being components of the nervous system. The cysteine arrangement of Aeq5-like1 and Aeq5-like2 had been reported in Aeq5 peptide extracted from total lysate of *A. equina*, where they were shown to be lethal for crabs and hence classified as a toxin. Neuronal localization of peptides of this type has not been reported before. Because multiple disulphide-rich venom components (including ShK toxin) are neurotoxins targeting neuroreceptors, the new molecules may potentially act through binding to endogenous receptors in *Nematostella*.

Toxin-like molecules have been reported in vertebrate and insect brains (43-46), but in many cases their sequence similarity to toxins is very low making convergent evolution a realistic scenario. One previously reported example is Agatoxin-like peptides in the insect neuroendocrine system where the sequence similarity to the corresponding spider toxin is quite high (45). The most characterized group of toxin-like neuroregulators are mammalian prototoxins with significant structural similarity to snake three-fingered toxins (43). Members of these family can be secreted or GPI-anchored (which makes them similar to the membrane-bound Aeq5-like2). They regulate function of nicotinic acetylcholine receptors at multiple levels, including direct binding to the mature receptor leading to modulation or inhibition of its activity as well as by affecting subunit stoichiometry during receptor assembly.

### Regulation of ShK-like2 by alternative promoters and transcription start sites

ShK-like2 is regulated by alternative promoters resulting in different dynamics of the transcripts encoding the two precursor isoforms in development and probably in tissues. As our exon usage and ISH analyses showed, ShK-like2b isoform is preferentially expressed during the first five days of development with initial expression in the ectoderm of early planula. ShK-like2a was first expressed in the late planula and mostly in the endoderm. Additionally, qPCR analysis suggests that the two isoforms might be expressed at different levels in the mesentery of adult females. Thus, ShK-like2 gene is equipped with a sophisticated regulatory system allowing expression flexibility.

However, ShK-like2a and 2b expression patterns largely overlap in primary polyps. Eukaryotic promoters not only initiate transcription but also affect mRNA stability, translation efficiency and subcellular localization (47, 48), particularly in neurons (49). Thus, even though ShK-like2a and 2b are co-expressed in the same cells, the regulation by different promoters may potentially result in different abundance and localization of protein precursors. Additionally, different promoters might respond differently to stimuli. Regulation by alternative promoters and TSSs is common in development (47, 50) and has been reported for multiple neuronal proteins, such as signaling molecules (51), receptors (52) and enzymes (53).

Alternative TSSs result in different preproparts in the ShK-like2 isoforms. Propeptides are known to affect the fate of mature peptide in multiple ways including its stability and localization (54). Different propeptide sequences may potentially affect sorting of mature ShK-like2 and lead to different intracellular localization of the two isoforms. For example, in the mammalian neuroendocrine system, prohormones are sorted into regulated secretory pathway through binding of their propeptide to a specific receptor in the *trans*-Golgi network (55) and therefore different propeptide sequences might affect the binding. Furthermore, in the sea slug *Aplysia californica*, neuropeptides derived from processing of the same precursor are sorted to different classes of secretory vesicles and transported to different neural processes of the same cell depending on the sequences of intermediates during the stepwise maturation (56).

Thus, the structure of the ShK-like2 gene underlies regulatory flexibility at the transcriptional level in development and may potentially affect regulation at post-transcriptional, translational and post-translational levels. The conservation of such gene structure among ShK-like2 homologs among anthozoans supports its functional importance. To the best of our knowledge, this is the first report of gene regulation by alternative promoters and TSS in a cnidarian and, in particular, in cnidarian neuropeptides.

## Methods

### 1. Sequence search and analysis

Toxin sequences deposited in UniProt (ToxProt database) were used as a query in BLAST to search the Transcriptome Shotgun Assembly (TSA) databases on NCBI using the online BLAST (https://blast.ncbi.nlm.nih.gov/Blast.cgi) with tBlastn option. The TSA dataset was also downloaded to screen for ubiquitous ShK cysteine domains using a regular expression screening approach. All of the TSA sequences for *Nematostella* were translated into their predicted proteins using Transdecoder (62). Predicted proteins were then screened using the unix command `grep` for the consistent amino acid arrangement of the last three cysteine residues: CXXXCXXC followed by either the stop codon (`C\w\w\wC\w\wC\*`), one amino acid residue prior to the stop codon (`C\w\w\wC\w\wC\w\*`), or two amino acid residues prior to the stop codon (`C\w\w\wC\w\wC\w\w\*`). Sequences with repetitive ShK domains were removed and the remaining predicted proteins were then screened with SignalP online tool (http://www.cbs.dtu.dk/services/SignalP/) (24). The remaining candidates were screened by eye and those with more than six cysteine residues in the predicted mature peptide or a longer or shorter than expected propart region were removed. To align protein sequences, Clustal Omega (57) (https://www.ebi.ac.uk/Tools/msa/clustalo/) and Muscle algorithm implemented in MegaX (58) were used. Gene expression data were retrieved from NvERTx database (http://ircan.unice.fr/ER/ER_plotter/home) (27).

The sequences from other sea anemones were retrieved using tBlastn on Reef Genomics (http://reefgenomics.org/) and NCBI TSA. To resolve isoforms of ShK-like2 orthologue in *A. equina*, a de novo transcriptome was assembled using Trinity (v2.6.6) (59) from previously published raw reads (accession SRR7507858 (36)). Assembled contigs were used as Blastx queries against the translated ShK-like2 sequences from *Nematostella*. After both ShK-like2a and b were identified we resolved their genomic loci and the exon-intron structure by performing Blastn against the *A. equina* genome on Reef Genomics.

### 2. Transgenic construct and immunostaining

The ShK-like2 gene fragment (scaffold_260:204,034-206,332; coordinates according to (30)), which includes putative promoter B, exon 2, intron 2, and exon 3, was amplified from genomic DNA. The ShK-like2 gene fragment containing exons 3 and 4 was amplified from *Nematostella* cDNA. A short sequence encoding the GGSGGSGRA peptide was introduced to the 3′ end. A mutation (underlined) was introduced into exon 3 in both fragments to generate a unique *Eco*RI restriction site (in bold): CAGAA<u>AAA</u>AG → CA**GAA**TTCAG. The two fragments were fused by ligation following restriction at the *Eco*RI site. Thus, a DNA fragment encoding full-length ShK-like2b under control of the presumed regulatory regions was obtained. It was subsequently cloned into the plasmid generated in our previous work (9) to be located upstream of the mOrange2 sequence by replacing the Nep3 regulatory sequence. This resulted in a construct encoding ShK-like2b connected to fluorescent reporter protein mOrange2 by a flexible GGSGGSGRA linker and under control of ShK-like2b regulatory sequence.

The transgenic plasmid was injected into *Nematostella* zygotes along with the yeast meganuclease I-*Sce*I (New England Biolabs, Ipswich, MA) to facilitate genomic integration according to an established protocol (60). Immunostaining was then performed using a commercially-available rabbit polyclonal antibody against mCherry (Abcam) diluted to 1:400, consistent with a previously published protocol (61). Imaging was performed with an Olympus FV-1000 inverted confocal microscope (Olympus, Tokyo, Japan) equipped with a 561 nm laser at the Advanced Imaging Unit of the Hebrew University. Images were processed with ImageJ to create Z-stacks of maximum intensity projections; contrast and brightness were adjusted manually.

### 3. ISH and dFISH

ISH and dFISH were performed as previously described (3, 7, 62) with addition of T1 RNAse treatment (2 u/µl, Thermo Fisher Scientific, Waltham, MA, USA) after probe washing in embryos older than 4 dpf to reduce background staining. Probe sequences are available in **Suppl table 2**. In dFISH, tyramide conjugated to Dylight 488 and Dylight 550 fluorescent dyes (Thermo Fisher Scientific) was used for visualization. Stained animals were visualized with an Eclipse Ni-U microscope equipped with a DS-Ri2 camera and an Elements BR software (Nikon, Tokyo, Japan). Photoshop (Adobe, San Jose, CA, USA) was used to manually adjust brightness and contrast levels of the images and to overlay different channels.

### 4. qPCR and RNA-seq

Total RNA from *Nematostella* larvae (2 dpf, 4 dpf, 6 dpf, 10 dpf), adults (males, females) and female body parts (tentacles, physa, and mesentery) was extracted as described in our previous work (9). cDNA was synthesized using SuperScript III Reverse Transcriptase (Thermo Fisher Scientific).

Using Primer Express Software (Thermo Fisher Scientific), we designed qPCR primers specific to the ShK-like2a and ShK-like2b transcripts (**Suppl table 3**). The forward primers map to exon 1 or exon 2 of ShK-like2 gene, for ShK-like2a and ShK-like2b respectively. The reverse primers map to boundaries between exon 1 - exon 3 or exon 2 - exon 3 for ShK-like2a and ShK-like2b, respectively. To quantify transcripts, SYBR(tm) Green PCR Master Mix (Thermo Fisher Scientific) and StepOnePlus Real-Time PCR System (ABI instrument, Thermo Fisher Scientific) were used. PCR amplification was carried out with equal amounts of the cDNA in triplicates. The HKG4 gene, whose expression was found very stable across *Nematostella* life stages (9), was used for normalization. ANOVA was used to determine if the expression of ShK-like2a and ShK-like2b was significantly different across development and tissue. Ct values were used to calculate relative abundance X of the transcripts using formula X=2^-Ct(transcript)^/2^-Ct(HKG4)^.

To test whether ShK-like2 undergoes alternative isoform expression across development in *Nematostella* we performed DEXseq analysis. An updated transcriptome for *Nematostella* was *de novo* assembled using Trinity (v2.6.6) (59) using previously published raw reads (PRJEB13676) (63). The transcriptome contained both ShK-like2a and ShK-like2b as isoforms of the same gene. SuperTranscripts were constructed by collapsing unique and common sequence regions among isoforms using the Trinity superTranscripts pipeline (59, 64). *Nematostella* raw reads (27) generated across development (1-8 dpf and juveniles with two replicates) were downloaded from NCBI sequence read archive (PRJNA418421 and PRJNA419631) and mapped to the superTranscripts assembly using STAR (65). DEXseq was performed to detect differential exon usage (32, 64).

### 5. Recombinant expression and purification

Fragments encoding ShK-like1 and ShK-like2 mature peptides were amplified from *Nematostella* cDNA and cloned into pET40 vector (MilliporeSigma, Burlington, MA USA) with deleted DsbC signal peptide sequence (9). The resulting constructs coded for fusion proteins DsbC-6×His-ShK-like1 and DsbC-6×His-ShK-like2. A TEV protease cleavage site (ENLYFQ/G) was introduced immediately upstream of the mature peptides; that resulted into addition of a Gly residue to N-terminus of ShK-like1 mature sequence. Heterologous expression was carried out in BL21(DE3) *E. coli* (MilliporeSigma) strain at 22°C overnight.

Nickel affinity FPLC purification of the fusion proteins was followed by cleavage with TEV protease. Reverse phase FPLC on a Resource RPC column (GE Healthcare, Danderyd, Sweden) in acetonitrile gradient in 0.1% trifluoroacetic acid was used for final purification of the mature peptides. The peptides were vacuum dried and stored at −80°C until use.

### 6. Toxicity assay

Zebrafish (*Danio rerio*) larvae younger than 120 hpf were kindly provided by Dr. Adi Inbal (The Hebrew University Medical School). At this developmental stage, any ethical permits are not required for experimental use according to the European and Israeli laws. ShK-like1 and ShK-like2 recombinant peptides were added to 0.5 mg/ml concentration to zebrafish media and incubated with 4 dpf larvae in 500 µl wells. Experiments were performed in duplicates. 5 mg/ml BSA was used as a negative control. Behavior and survival were recorded under an SMZ18 (Nikon) stereomicroscope.

### 7. Analysis of nematocyst content

Nematocysts were purified from 4 dpf planulae using a Percoll gradient based on a protocol kindly provided by Monterey Bay Labs Ltd (Caesaria, Israel). Briefly, the planulae were frozen and homogenized in 0.01% Triton X100 in 12.5 ppt ASW. Cell debris was removed by two centrifugation steps (2500 × *g* and 2000 × *g*). The pellet was re-suspended in 12.5‰ ASW and laid on top of 80% Percoll (MilliporeSigma) in 12.5‰ ASW. Centrifugation at 1500 × *g* for 10 min resulted in nematocyst pellet which was re-suspended in 6‰ ASW followed by exchange to 1‰ ASW.

The purified nematocysts were disrupted by ultrasonication, centrifuged (24000 × *g*, 10 min) and the supernatant was sent to LC-MS/MS analysis. MS analysis was performed using a Q Exactive Plus mass spectrometer (Thermo Fisher Scientific) coupled on-line to a nanoflow UHPLC instrument (Ultimate 3000 Dionex, Thermo Fisher Scientific) at the Proteomics Center of the Alexander Silberman Life Sciences Institute, The Hebrew University of Jerusalem. Protein identification was performed using *Nematostella* protein models (David Fredman; Michaela Schwaiger; Fabian Rentzsch; Ulrich Technau [2013]: *Nematostella vectensis* transcriptome and gene models v2.0. figshare. Data set; https://doi.org/10.6084/m9.figshare.807696.v1) with manually added ShK-like1 sequence and MaxQuant software (66) as described earlier (9). The raw LC-MS/MS files along with MaxQuant output were submitted to ProteomeXchange Consortium via the PRIDE (67) partner repository with identifier PXD019085.

To rank protein abundancies, intensity value was normalized against the number of unique and razor peptides identified for each protein group. Among the top 30 proteins, uncharacterized proteins were annotated using InterProScan (https://www.ebi.ac.uk/interpro/) search for protein domains and the SignalP tool to identify secreted proteins.

### 8. ShK-like1 and ShK-like2 activity on polyps

Eleven days old primary polyps were placed into a 96 well plate in the number of 10 – 14 individuals/plate to minimize physical contact between them. Recombinant peptides were added to the concentration of 0.5 mg/ml in the final volume of 90 µl/well. BSA (5 mg/ml) and NvePtx1 toxin (9) were used for comparison. The plate was placed under an SMZ18 (Nikon) stereomicroscope and effects were recorded in two wells simultaneously (ShK-like2 - BSA; ShK-like2 - NvePtx1; ShK-like2 - ShK-like1) every 30 minutes during 2 h avoiding any physical disturbance of the animals.

### 9. Knockdown with Morpholino oligonucleotides

Morpholino antisense oligonucleotides (**MO; Suppl table 4**) were ordered from Gene Tools, LLC (Philomath, OR, USA). MOs were designed to complement sequences corresponding to the boundaries of exon 3 – intron 3 (E3-I3) and intron 3 – exon 4 (I3-E4) of ShK-like2 gene to interfere with splicing. MOs were injected into *Nematostella* zygotes at the concentration of 0.9 mM in parallel to a control MO, which has no binding target in *Nematostella*. The embryos were maintained in 16‰ ASW at 22°C in the dark for 10 days and their development was monitored daily. After 10 days, the proportion of polyps that were unsettled vs. settled was recorded, and Fischer’s exact test and Chi-square test used to calculate significance.

To confirm that MOs affected splicing either by provoking exon skip or intron retention, RNA was extracted from the 2 dpf embryos, cDNA was synthesized and targeted exon – intron boundaries ShK-like2 gene were amplified by PCR with specific primers (**Suppl table 3**).

## Acknowledgments

We would like to thank the following researchers of the core facilities of the Alexander Silberman Institute of Life Sciences, The Hebrew University: Dr. William Breuer (Interdepartmental Unit) for the help with mass spectrometry, Dr Mario Lebendiker (Protein Expression and Purification Unit) for the help with chromatography, and Dr Naomi Melamed-Book (Advanced Imaging Unit) for her help with confocal microscopy. This research was supported by an Israel Science Foundation grant 869/18 to YM, U.S.-Israel Binational Science Foundation grant 2014667 (NSF Award 1536530) to YM and AMR.

## Notes

### Competing Interest Statement

The authors have declared no competing interest.

