## Supplementary tables for "Toxin-like neuropeptides in the sea anemone *Nematostella* unravel recruitment from the nervous system to venom"

**Suppl table1.** Annotation of the top 30 most abundant proteins identified by LC-MS/MS of the soluble content of planulae nematocysts. The uncharacterized proteins annotated by InterProScan are **in bold**.

| Annotation | Gene model | Normalized intensity | Signal peptide | Reference |
|---|---|---|---|---|
| **Ig-like dom** | NVE12991 | 1,24E+08 | - | |
| Nep4 | | 1,10E+08 | + | [7] |
| Nep6 | | 8,21E+07 | + | [7] |
| Cys-rich | NVE21061 | 7,49E+07 | + | |
| Nep3-like | | 5,56E+07 | + | [9] |
| NR2 | NVE3844 | 5,48E+07 | + | |
| Shk-like1 | - | 5,36E+07 | + | |
| Nematogalectin | NVE3845 | 3,30E+07 | + | |
| **LamG-like dom** | NVE25957 | 2,58E+07 | + | |
| **CAP+C-lectin-like dom** | NVE17241 | 2,45E+07 | - | |
| Nep3 | | 2,06E+07 | + | [7] |
| **WAP dom** | NVE11707 | 1,87E+07 | + | |
| **VWF_A dom** | NVE3052 | 1,75E+07 | + | |
| **CRISP-related 1** | NVE531 | 1,75E+07 | + | |
| Unknown 1 | NVE900 | 1,75E+07 | + | |
| Nep16-like | NVE21616 | 1,72E+07 | + | |
| **EGF-like dom** | NVE12020 | 1,48E+07 | - | |
| Nematogalectin homol | NVE3843 | 1,48E+07 | + | [39] |
| Nep8-like | | 1,41E+07 | + | |
| **CRISP-related 2** | NVE9188 | 1,22E+07 | + | |
| **DKK-like 1** | NVE22167 | 1,21E+07 | + | |
| Unknown 2 | NVE26126;NVE2447 | 1,18E+07 | - ; + | |
| Prolyl isomerase | NVE9332 | 1,18E+07 | - | |
| **Histone** | NVE22351;NVE19301;NVE8849;NVE8840;NVE6466;NVE5231;NVE5226;NVE2470;NVE24648;NVE24630;NVE24360;NVE24358;NVE24175;NVE24050;NVE24024;NVE24019;NVE24004;NVE23384;NVE23184;NVE22196;NVE21356;NVE21353;NVE20560;NVE19440;NVE18669;NVE14707;NVE14104;NVE22353;NVE24230;NVE2880;NVE24857 | 1,10E+07 | - | |
| **Lectin** | NVE2180 | 1,06E+07 | - | |
| **γ-glutamyltransferase** | NVE20501 gamma-glutamyltransferase 1 | 1,03E+07 | - | |
| Nep16 | Nep16 | 9,95E+06 | + | [7] |
| GAPDH | NVE23813 | 9,56E+06 | - | |
| **DKK-like 2** | NVE144 | 9,49E+06 | + | |
| **MACPF dom** | NVE160;NVE2369 MAC/Perforin domain; - | 9,20E+06 | + ; - | |

**Suppl table 2. Sequences of ISH and dFISH probes (sense).** The sequence shared between ShK-like2a and ShK-like2b isoforms is in bold.

| Transcript | Sequence (sense) |
|---|---|
| ShK-like1 | taatcagtcacattgaatgaggctcatattcccaaacattactggctgaaaataccct<br>aaaatgccagttatttgctttaattatttaccagtggatactcccacatagtgggaca<br>accataagaaccaatgcatgaagaatagccttataaaggtcgctcatataatatctg<br>ccataacatttggtgaacaacttaaaggaccaggatctcagctcgtttaacatggccc<br>gaaagctcctggcagtgctaatggtgtgtaccttttcctaatcgctgcctcgatggg<br>aaccaatgccctccctttccatgagggaatagagcgtcgagcagccaaatgtgttgac<br>aagatgccgtttgtgtgcatgcgaaaagatatcccggccatttgtaaaaatcgaaatc<br>atcgaagttatgccttcattatggacgtgtgccgcaaaacatgcggtcagtgcaccta<br>agcaagaacggctgtatttcgagctgaaacgcaggctaccagcgatggagtcacgaat<br>gaataaatttgagtaaccaaccaaaccctctgttgtgagatttattgg |
| ShK-like2b-long | actcatccgtctctacattcgcgagtcacagctgtgaggagagccaccttttagccaa<br>aacacagtcaactctaactcgccatggacgtcaaattggttgcaatccttttcgcatg<br>cacgttgttctcgctctcgttcgctaatggatatagactagaaagcttgagaaatctg<br>ccaaatgacgctttggaggaggagagtgag**gttgcagaaaaaagcccactgaagaagc<br>gaggttgctctgatgcgtttcctgttgtttgccggtccccaagcgtgaaagctgcttg<br>ttataaccctaaccataggagccacgccttcataaccgacgtctgcaagcacacgtgc<br>catctttgttaaatacgggaaatgactgcgatggagataaagatgacgagaaacaaga<br>atacataatcaaataaaatgagcccagaacgttgtacaaaaacctcccctcccttatg<br>tattcagtgtagtatagcatagtgtattacaatctccagtgtagtgctcccttcgctc<br>ctctatataccataagattgtacaggtgtgaacatataataaagatgtgtcagatgaa<br>tcagtcgaataaagactgggaaa** |
| Shk-like2a-short | cgtgatataatagtcccttattgccctggtagcataataactgacactgatagtcgcg<br>tcggtcaaagcatcaaaagggaagggtataaaacgaagctgcgttggcgcaacatcca<br>aacagtgcacacaagcaaacaagaaggtcagaccctctgctaacagagctacaaaaaa<br>aacacaagagaaaaatgacgaccattcgatgcgtgcttttcgctgtattactgtttgc<br>ttattgcgctttgttgataaaagcgcgctcgattgatgctgaggccgagaagacctgg<br>caagaggaggagacaaaaaca**gttgcagaaaaaagcccactgaagaagcgaggttgct<br>ctgatgcgtttcctgttgtttgcc** |
| ShK-like2b-short | actcatccgtctctacattcgcgagtcacagctgtgaggagagccaccttttagccaa<br>aacacagtcaactctaactcgccatggacgtcaaattggttgcaatccttttcgcatg<br>cacgttgttctcgctctcgttcgctaatggatatagactagaaagcttgagaaatctg<br>ccaaatgacgctttggaggaggagagtgag**gttgcagaaaaaagcccactgaagaagc<br>gaggttgctctgatgcgtttcctgttgtttgccggtccccaagcgtgaaagctgcttg<br>ttataaccctaaccataggagccacgccttcataaccgacgt** |
| Class8-like | atttgaacgttcaactaacaagcgtcttagtctcctgacaagaaaaactctctatctc<br>tcaaaatgcgtactctggtggttctcctcatcggcgctgtcctcctctgctctgccaa<br>tgctttcctcgacgagcttctggccgagagcgtgaacgacatgacagacaagcgtgca<br>tgcttcgacaaatacaagtccaacatctgtggtggtgtcatcagccccgctcactgcg<br>tgaggaggagtggtcgcatggccaagttcgcaaaggagaactgtgcgcacttttgtgg<br>attctgttagggaaaacagtgatgtgaagacgaagacgaaatggaaccggctggactg<br>gctttcacctcgacagattatgattgttattatagggcccttatagttccatataata<br>aatagttgaaaagcagaa |
| Aeq5-like1 | catttgtacggatcacgcctagctacgcgaagccgtttgataaataaaaggcaagaaa<br>tacttttacaaagatgaagtccgtaattgcagttcttgttctctcgctggttctcgtc<br>aacttcacacaagcagcgaaggacgatcgatggaaggcgtgccgcatgaagtgctaca<br>ccgaaagcaaactgtgcatgaacaacgactccaagtgtttcgactcccaatcgtgtaa |

| | |
|---|---|
| | tagctgcattcaacaagtatacagcccgtgtttcaatagatgccaagagatgctccga<br>cgacgagaggcctttaaacgtatgttcgcatttgacgaggagaattaaagaccaagaa<br>ctatattccggacattcaattgtttttatctaaaacactgtaatgatacaccaccaag<br>ttctgcaaagacatacatcggaaacctatatatttagataaaatttcaacactcattt<br>ttggtctatgagtttttttgacctgtttataagctggtcaactttgcacccacgtta<br>ttttacgtaacggccaggaagtttctcatttgtagaaactagtaacggcaaatcacaa<br>cttgtgactgaatcacgtggttagatagggagaggagaggaaggagaatcgccttcgt<br>tcacttagataca |
| Aeq5-like2 | atcgggatcgtagtggctcagtgccgtcgggacttggagtcaaaccgtacccttgatt<br>atcgcactaggatgaggacttcattaattcttgttgccatggtgatggtgagtgttct<br>gctcccgtacacctacggctcatcatgtgactccttctgtaccgagcaagcgaacaag<br>tgtctgacgggctgtgagggcttcgtggggtgtatggagtgcacgaacttcgccggcc<br>actgtcgggagcaatgtcgaaagagatccgtcaagagacgcaaggagattcgagcgcg<br>atttacaaaagaacccacagaagaatcttaaccaccacccccctcccccctgggtttgat<br>gacgtcacaagaaattacatggactacgtaaaccattgcttttaagcgacaaatcata<br>ttactttgtctttgtgtaaatgcatactatatggtaataccctcaattgtaaagatat<br>ttgcc |
| ShK-like3 | TCGTTGAAACGGAATCCGCCAAACAAACCTCTGGCGCCCGGATACAGTATAAAGCGATCTACGT<br>GTGCTCTAAACTATAGGCAACATTTATTCTCCACTGACTCAAGTGTGGCTGATGCCGAACGCGT<br>CACACAAATAGGAAGTCATGTACCGAAAGCTTGCCATCGCGGTCCTCCTGTGTTCTATCCTTTT<br>CTCTGGAGGTCTGGGCTCTAATGCATCAAAGGAAGTAAAGGTAACTGCTAAAAAGAAAACTGCT<br>TTGCTTTGCTTCGATGCATACCCATCCCTTTGTCGTAAATCGGTAGCCAGGAAGGCATGCAATA<br>ATTCAGATCACCGTAGCCATGCGTTTGTGATCGACGTCTGCAGGAAAACGTGTGGTCGCTGCTA<br>AAACAATGTGGGACATATGAAAGCCAAGTTTCACAATTAGGCAAAAACGTACAGTGAACAGTGA<br>AGAAACGGCA |
| ELAV1 | cttggtaccgagctcggatccactagtaacggccgccagtgtgctggaattcgccctt<br>gacgttcagttacgctcacggcttatttcagttcatttttggtgaccccgtctctaat<br>ttattcgttcgattttctcagctcagtgaaactgagacactttctgtgatgttggaca<br>tgatggacgacaattctaaggctatggctgaagatatcaacaatcacgaaccgatgga<br>aaatggaacttcagatgaaagaacaaatcttataatcaactacgtgcctccgagtatg<br>agccaagaggatatcaaaaaaatatttgggactgtgggaaatgtcacaagttgtaaac<br>tcatccgcgatcgagcaacaggacagagccttggctacgcgttcgtaaactacgataa<br>tcctgacgatgcgaacaaagctgtaagggagatgaatggagctcgacttcaaaataaa<br>accctgaaagtgagcttcgcgcgaccttcttctaccgaaattaaaaacgcgaatttat<br>atatcagcggtctaccgaaagacatgaaagaagaagaggtcgaagcattgtttaagcc<br>gttcggaaaaatcataacttctaaagttctgaaagatgtgagcggcgaaggtaggggc<br>acaggatttgtacgttttgacaagcgctgtgaagctcaaacggccattgatgacctga<br>ataataaaacattacccggcactaatgttaaactcacagtaaagttcgcaaatccacc<br>gaattcaagacagccagctatgccgcttagtccggcgctaaccagccctctaggaaga<br>gcgctaactccccagcgaaacttcagtggagggccggttcatcatcagatgttgaaca<br>tgaggtattcccctatgacagcgagtagcttctcccctgcttgccaagccaattgaag<br>ggcgaattctgcagatat |

### Suppl table 3. Primer sequences

| | Transcript | Sequence |
|---|---|---|
| Splicing effects of MOs | Exon2_foward | CGCTCTCGTTCGCTAATGG |
| | Intron3_reverse | GCTCTCTATTGCGAATAAGATAGCG |
| | Exon3_foward | GAAAAAAGCCCACTGAAGAAGCGAG |
| | Exon5_reverse | CAATCTTATGGTATATAGAGGAGCG |
| | ShK-like2a_foward | TGATGCTGAGGCCGAGAAG |

| | | |
|---|---|---|
| qPCR | ShK-like2a_reverse | GGGCTTTTTTCTGCAACTGTTT |
| | ShK-like2b_foward | CGCTCTCGTTCGCTAATGG |
| | ShK-like2b_reverse | GGCTTTTTTCTGCAACCTCACT |
| | HKG4_foward | CTGCCAAGAAGAATGCAGCTG |
| | HKG4_reverse | CCTTGAGTACCTTAGGCTGATGG |
| Genomic DNA sequencing | ShK-like1_foward | AAAGCTCCTGGCAGTGCTAATGG |
| | ShK-like1_reverse | GGCACACGTCCATAATGAAGGC |

**Supplementary table 4. Morpholino oligonucleotide sequences**

| MO | Sequence |
|---|---|
| E3-I3 | AAAGAGGCGTTGAACTTACCTCGCT |
| I3-E4 | ACCTGAGAAAGGAGGTACAGACCGA |
| Control | CCTCTTACCTCAGTTACAATTTATA |

**Supplementary table 5. Results of E3I3 MO injections in parallel to the Control MO**.

| | Replicate 1 | | Replicate 2 | | Replicate 3 | | Replicate 4 | |
|---|---|---|---|---|---|---|---|---|
| | E3I3 | Control | E3I3 | Control | E3I3 | Control | E3I3 | Control |
| Settled, # | 94 | 81 | 0 | 50 | 26 | 112 | 42 | 112 |
| Unsettled, # | 104 | 25 | 139 | 88 | 97 | 62 | 64 | 14 |
| Settled, % | 47.5 | 76.4 | 0 | 36.2 | 21.1 | 64.4 | 39.6 | 88.9 |
| Δsettled, % | -28.9 | | -36.2 | | -43.2 | | -49.3 | |
| Average Δsettled, % | -39.4 | | | | | | | |
| SD Δsettled, % | 8.8 | | | | | | | |

**Supplementary table 6. Results of I3E4 MO injections in parallel to the Control MO**.

| | Replicate 1 | | Replicate 2 | | Replicate 3 | |
|---|---|---|---|---|---|---|
| | I3E4 | Control | I3E4 | Control | I3E4 | Control |
| Settled, # | 0 | 224 | 0 | 142 | 0 | 123 |
| Unsettled, # | 107 | 14 | 95 | 44 | 56 | 31 |
| Settled, % | 0 | 94.1 | 0 | 76.3 | 0 | 79.9 |
| Δsettled, % | -94.1 | | -76.3 | | -79.9 | |
| Average Δsettled, % | -83.4 | | | | | |
| SD Δsettled, % | 9.4 | | | | | |
