## Supplementary figures for "Toxin-like neuropeptides in the sea anemone *Nematostella* unravel recruitment from the nervous system to venom"

**Suppl fig 1 Expression dynamics of the new genes at the transcriptomic (A) and proteomic (B) levels**

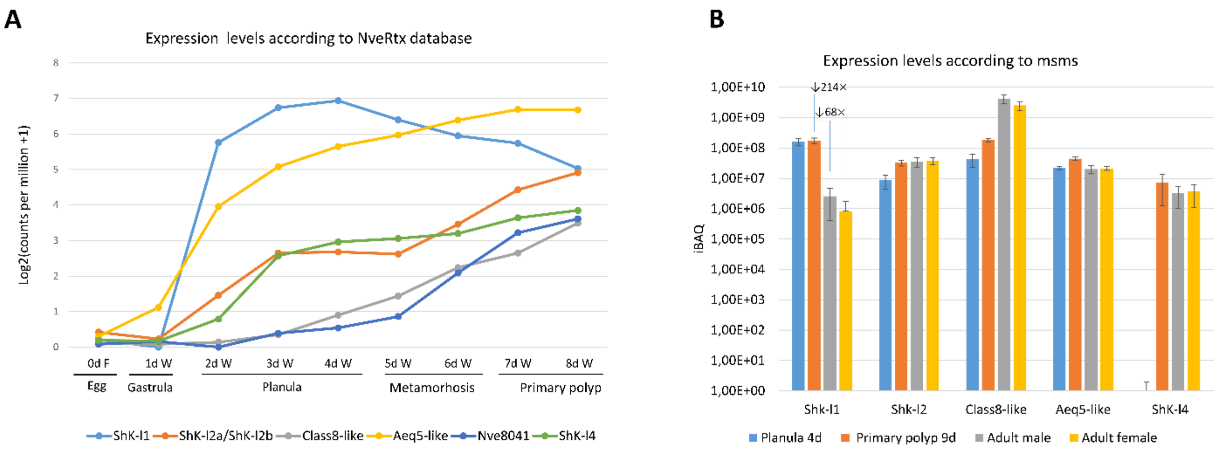


**Suppl fig 2. Co-localisation of ShK2a/2b by dFISH**

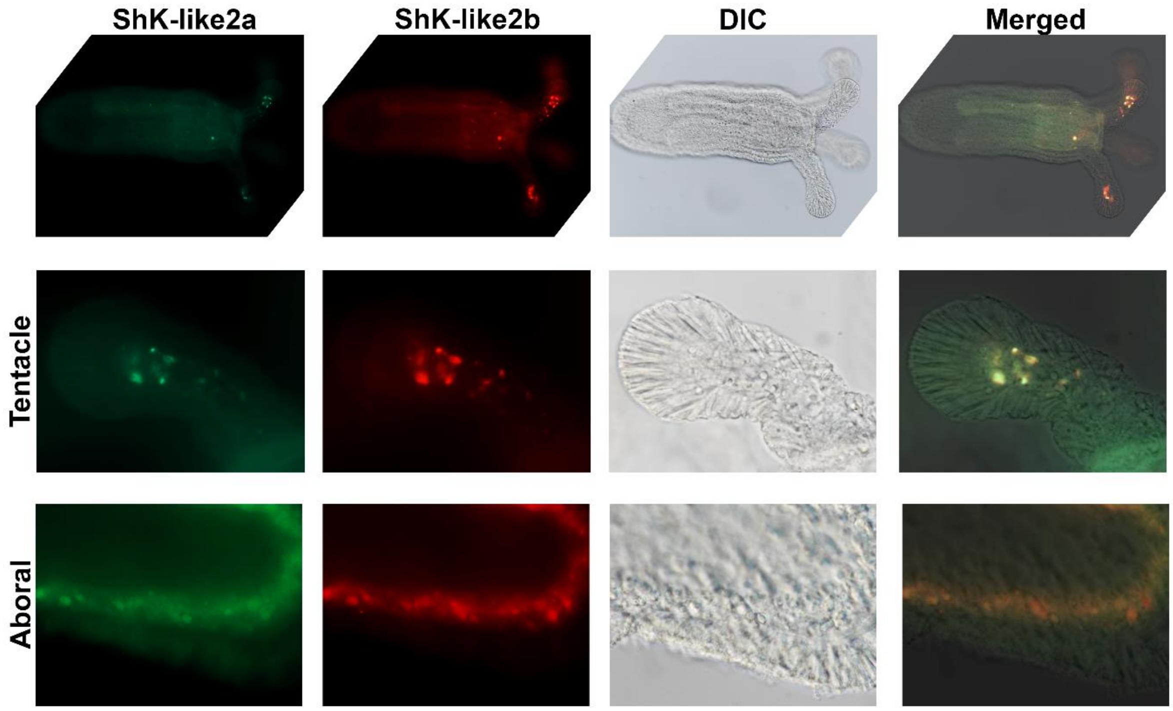

**Suppl fig 3. ISH of ShK-like3 in primary polyp.**

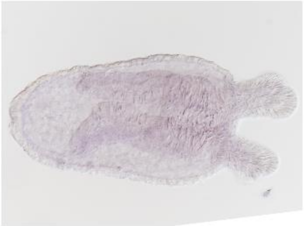

**Suppl fig 4. Co-localisation of ShK-like2 and ShK-like4 with ELAV expression by dISH**

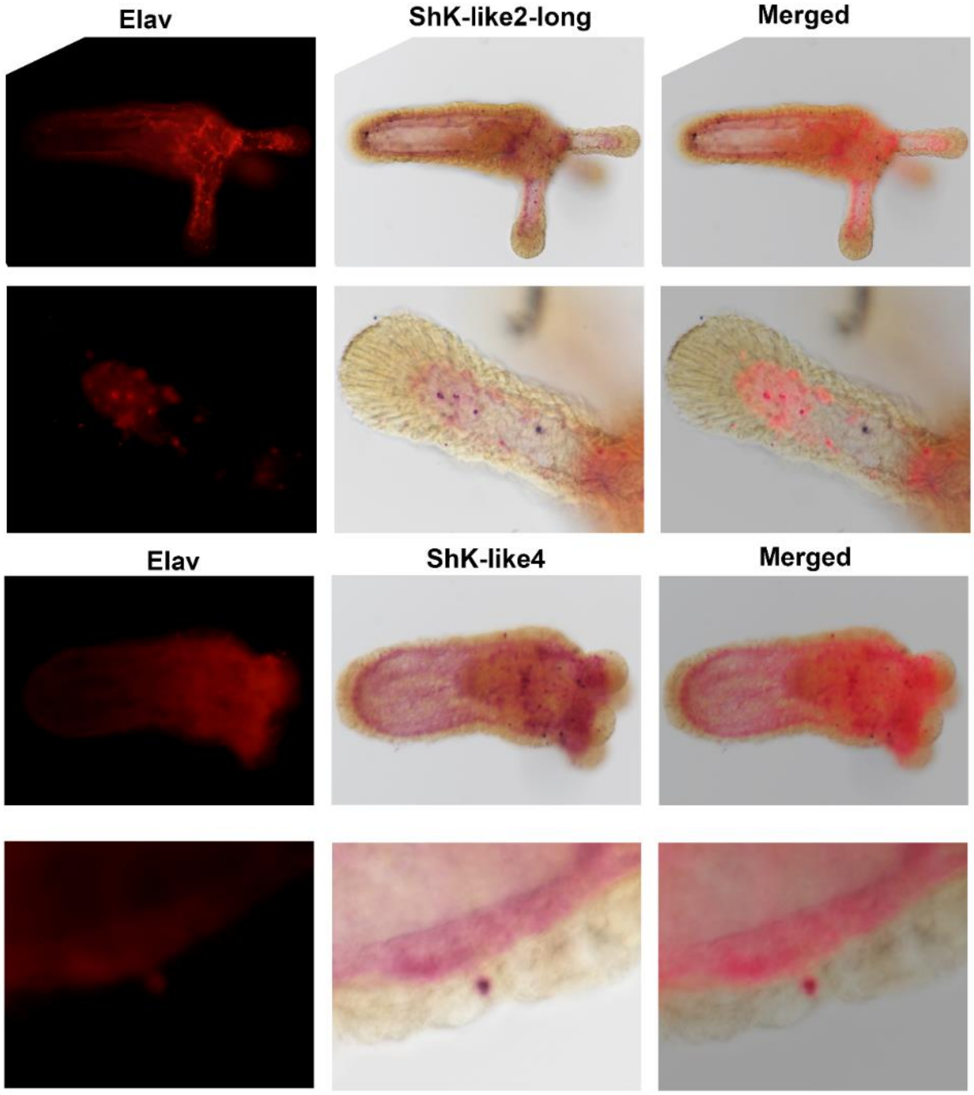

**Suppl fig 5. Co-localisation of Class8-like and Aeq5-like1 with ELAV expression by dISH**

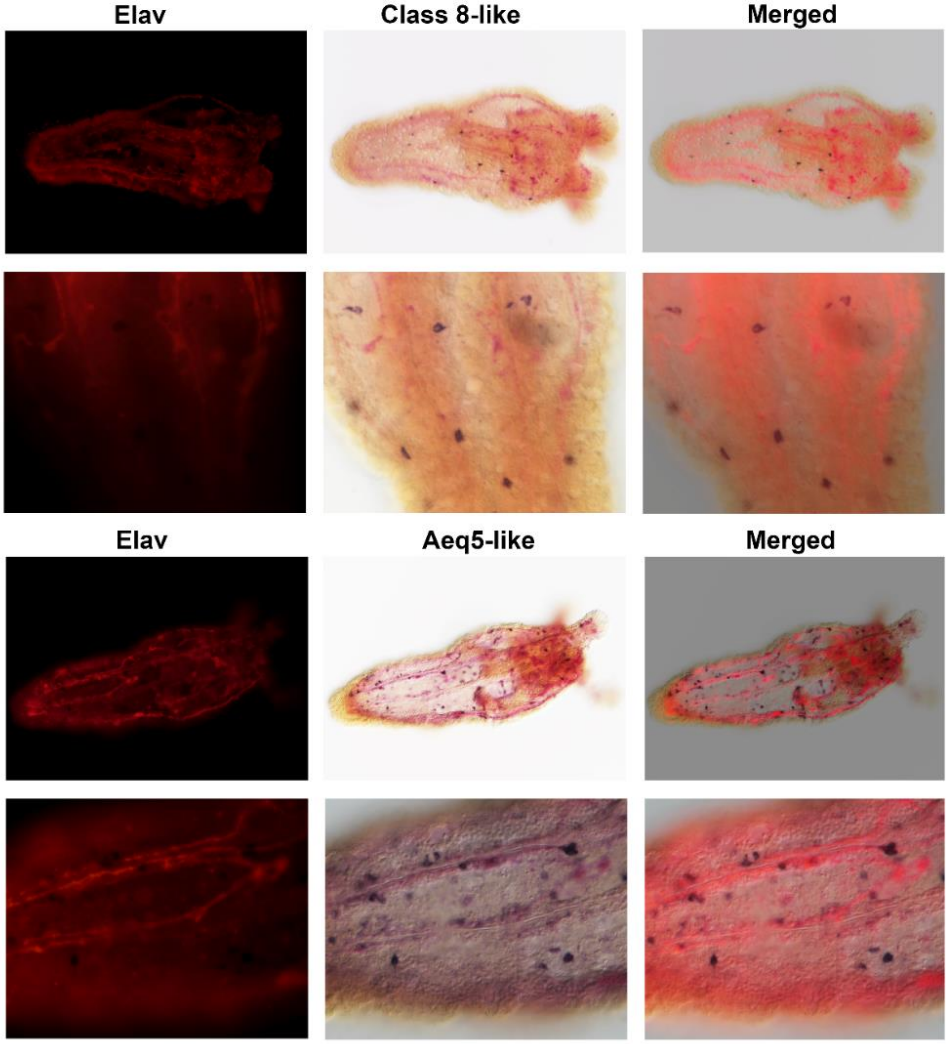


**Suppl fig 6. Expression pattern of the toxin-like neuropeptides according to the scRNAseq data**

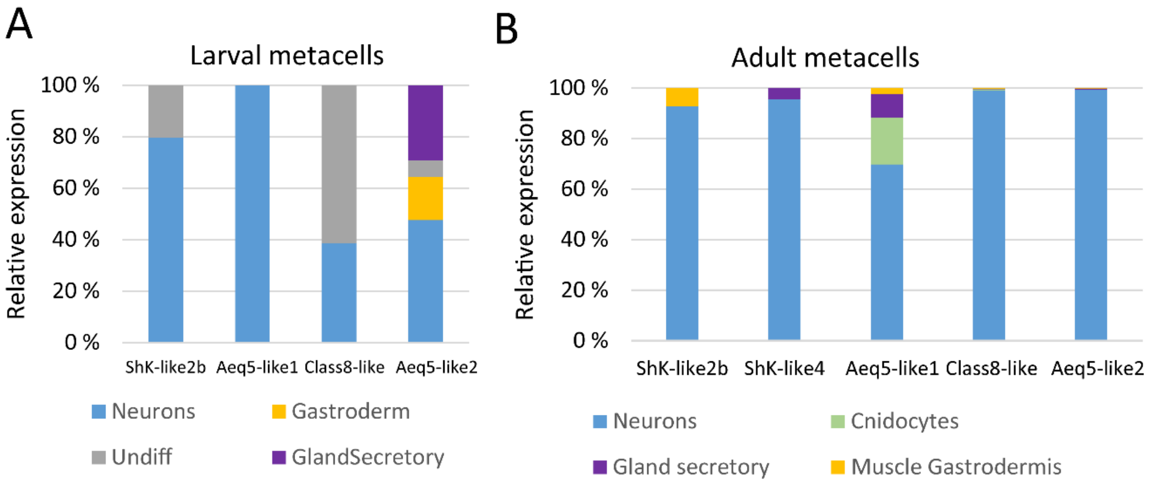

**Suppl fig 7. PCR analysis of ShK-like2 splice variants following E3I3 (A) and I3E4 (B) MO injections.** Yellow arrows point to fragments corresponding to disrupted splicing (intron 3 retention in A or skipped exon 4 in B); white arrows correspond to properly spliced ShK-like2 transcripts.

A. E3I3 MO injections

```
Possible products:
E2 to I3 retaining - 498bp
E2 TO E4 - 227bp
E2 TO E5 - 415bp
E2 TO E5 retaining I3 - 3578bp
E2 TO E4 retaining I3 - 3390bp
```

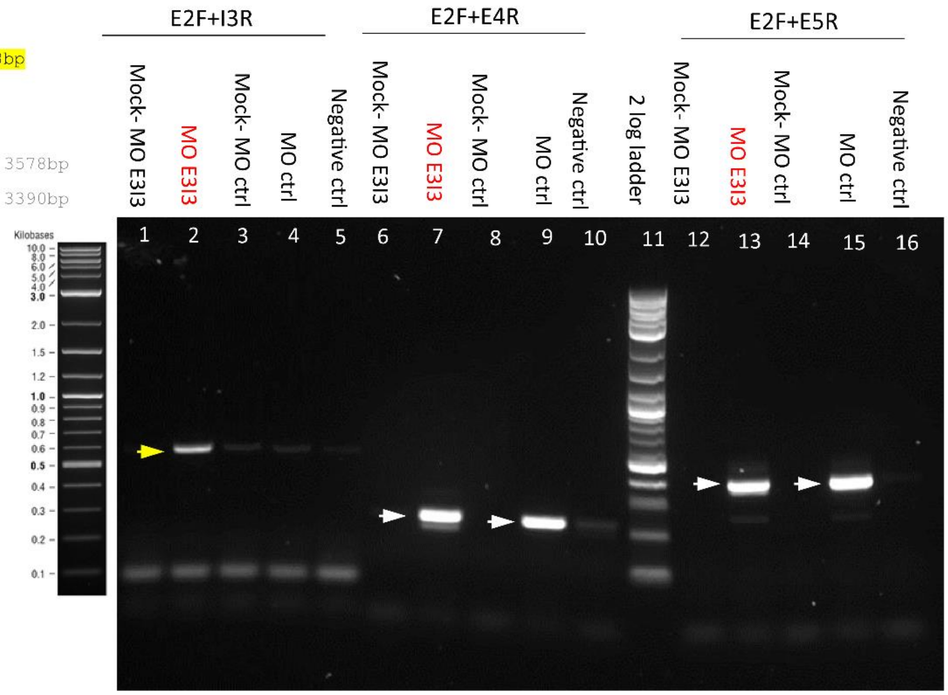


B. I3E4 MO injections

```
Possible products:
E2 to I3 retaining - 498bp
E3 TO E4 - 144bp
E3 TO E5- 337bp
E3 TO E5 skipping E4 - 201bp
E3 TO E5 retaining I3 - 3500BP
E3 TO E4 retaining I3 - 3307bp
```

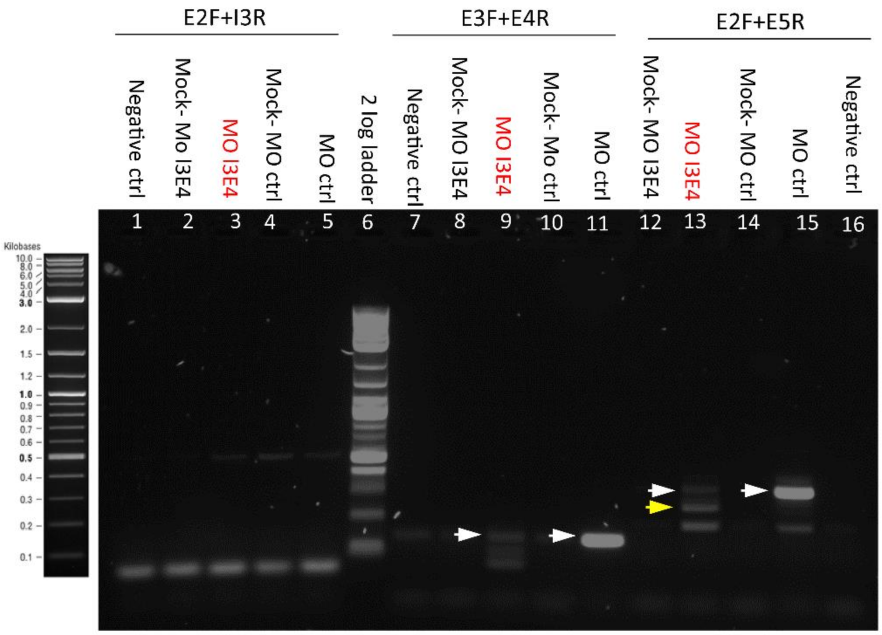
